# *Nicotiana benthamiana XYLEM CYSTEINE PROTEASE* genes facilitate tracheary element formation in interfamily grafting

**DOI:** 10.1101/2022.12.28.522093

**Authors:** Chaokun Huang, Ken-ichi Kurotani, Ryo Tabata, Nobutaka Mitsuda, Ryohei Sugita, Keitaro Tanoi, Michitaka Notaguchi

**Author notes:** **Corresponding Author:** Michitaka Notaguchi, Bioscience and Biotechnology Center, Nagoya University, Furo-cho, Chikusa-ku, Nagoya 464-8601, Japan. Material distribution footnote: “The author responsible for distribution of materials integral to the findings presented in this article in accordance with the policy described in the Instructions for Authors (https://academic.oup.com/plcell/pages/General-Instructions) is: Michitaka Notaguchi.”.

## Abstract

Grafting is a plant propagation technique widely used in agriculture. A recent discovery of the capability of interfamily grafting in *Nicotiana* has expanded the potential combinations of grafting. In this study, we showed that xylem connection is essential for the achievement of interfamily grafting and investigated the molecular basis of xylem formation at the graft junction. Transcriptome and gene network analyses revealed gene modules for tracheary element (TE) formation during grafting that include genes associated with xylem cell differentiation and immune response. The reliability of the drawn network was validated by examining the role of the *Nicotiana benthamiana XYLEM CYSTEINE PROTEASE (NbXCP)* genes in TE formation during interfamily grafting. Promoter activities of *NbXCP1* and *NbXCP2* genes were found in differentiating TE cells in the stem and callus tissues at the graft junction. Analysis of a *Nbxcp1;Nbxcp2* loss-of-function mutant indicated that *NbXCPs* control the timing of *de novo* TE formation at the graft junction. Moreover, grafts of the *NbXCP1* overexpressor increased the scion growth rate as well as the fruit size. Thus, we identified gene modules for TE formation at the graft boundary and demonstrated potential ways to enhance *Nicotiana* interfamily grafting.

## Introduction

Grafting is a technique used to unite two plant segments, following tissue adhesion and vascular reconnection. It has been widely practiced in horticulture, applied to propagate ornamental and fruit trees from ancient times. Moreover, grafting has been used for cultivation of vegetables in recent decades (Yamakawa, 1983; Kurata, 1992; Mudge et al., 2009; Goldschmidt et al., 2014; Aloni, 2021). A graft establishment depends on the successful formation of graft union that includes wound response, cell-cell adhesion, callus proliferation, and vascular formation (Melnyk, 2017; Rasool et al., 2020). Although there is a range in the extent of graft viability and hardiness, successful grafts achieve subsequent scion growth and/or fruit setting eventually. Previous observations clearly showed that vascular connection between the scion and stock is a determinant of grafting success (Proebsting, 1928; Kawaguchi et al., 2008). In general, graft compatibility is observed between phylogenetically closed relatives, limiting the range of plant combinations available for grafting. However, recent studies have shown potential possibilities to expand graft capability even among interfamily relations in eudicots and monocots (Notaguchi et al., 2020; Reeves et al., 2020). Subsequent studies showed that a part of Solanaceae and a part of parasitic plants in Orobanchaceae and Convolvulaceae exhibit tissue adhesion ability with interfamily partner (Kurotani et al., 2020; 2022; Okayasu et al, 2021). In *Nicotiana* interfamily grafting, new xylem connections were observed at the graft junction, as xylem bridges were seen in parasitism between interfamily host-parasite combinations (Cui et al., 2016; Wakatake et al., 2020; Kokla et al., 2022). However, phloem connections are not achieved in many cases of parasitism and are rarely established in *Nicotiana* interfamily grafting, whereas symplastic connections, probably through plasmodesmata, are established in both cases. The grafted and parasitic plants can survive and complete their life cycle until seed setting, implying essential roles of xylem bridge/connection in parasitism/grafting through these tissue adhesive conditions.

Tracheary elements (TEs), main components of xylem, are dead hollow cells with secondary cell walls that form vessels or tracheids (Fukuda, 1997; 2004; Fukuda and Ohashi-Ito, 2019). TEs are conducting tissues that transport water and dissolved minerals from the roots to shoots to maintain normal growth (Turner et al., 2007; Raven, 2013). In addition, proteins, hormones, and metabolites transported in the xylem have been evidenced to achieve various roles in squash, cucumber, cabbage, and maize plants (Satoh et al., 2006; Alvarez et al., 2008; Ligat et al., 2011). The *in vitro* xylem vessel element formation system revealed that certain plant-specific NAC-domain (no apical meristem, <u>N</u>AM; Arabidopsis transcription activation factor, <u>A</u>TAF; up-shaped Cotyledon 2, <u>C</u>UC2) transcription factors were identified as VND (<u>V</u>ascular-related <u>N</u>AC-<u>D</u>omain) genes (Kubo et al., 2005). It has been suggested that in Arabidopsis *VND6* and *VND7* are the primary determinants of TE differentiation and *VND1–VND5* are the weaker determinants (Kubo et al., 2005; Ohashi-Ito et al., 2010; Zhou et al., 2014; Endo et al., 2015). Moreover, *VND7*, a master switch in xylem vessel formation, has been demonstrated to regulate the expression of a broad range of genes in root and shoot (Yamaguchi et al., 2008; 2010; 2011). TEs undergo programmed cell death (PCD), removing the entire protoplast and forming a hollow vessel for substance transportation. Metacaspase 9 (*AtMC9*), *XCP1*, and *XCP2* degrade cellular remnants within the cell lumen after vacuolar rupture and assist in the final process of PCD (Funk et al., 2002; Avci et al., 2008; Bollhöner et al., 2013).

In this study, we investigated significance of xylem formation at the graft junction by two means; a comparison between successful and unsuccessful interfamily grafting and a comparison between with or without an auxin transport inhibitor, 2,3,5-triiodobenzonic acid (TIBA), that blocks xylem formation. Through the transcriptomic analysis, we have described the expression patterns of xylem-associated genes in successful and unsuccessful interfamily grafting. Using 36 transcriptome datasets and a super computer, a Bayesian network analysis was conducted to reveal the causal relationship between gene expression. Out of genes included in core modules for xylem formation, we focused on *NbXCP* genes to investigate how *de novo* TEs at the graft junction is important for the *N. benthamiana* interfamily grafting.

## Results

### Xylem reconnection is required for the scion growth

*Nicotiana benthamiana* grafted onto the inflorescence stem of Arabidopsis (*Nb*/*At*) constituted a successful interfamily graft combination. However, *Glycine max* and Arabidopsis (*Gm*/*At*) form an incompatible interfamily graft (Notaguchi et al., 2020). Xylem cells differentiated from *N. benthamiana* scions and attached to the cells in Arabidopsis stem tissues in *Nb*/*At* graft unions. We observed xylem bridges connecting the scion and stock at around 14 days after grafting (DAG). Fewer xylem cells differentiated from *G. max* scions in *Gm*/*At* graft unions. The xylem bridge was not formed even at 14 DAG (Figure 1A). In addition, we observed the development of secondary xylem in *N. benthamiana* and *G. max* stems from neighboring graft boundaries. Secondary xylem tissues were developed in *N. benthamiana* scion stems but were rarely observed in *G. max* scion stems (Supplemental Figure S1). Scion growth and fruit setting were observed in *Nb*/*At* grafts but scions did not grow in *Gm*/*At* grafts (Figure 1, B and C). In *Nb/At* interfamily grafting, tracheary elements (TEs), which have spiral patterns on cell walls (Figure 1D), were differentiated in the callus of the *N. benthamiana* scion. The *de novo* TEs were differentiated in the callus at the graft boundary and eventually connected the xylem tissues originally located in the scion and stock plants and formed xylem bridges between the scion and stock. Next, TE formation in *Nb*/*At* interfamily grafting was blocked to examine the effect of *de novo* TE formation on scion growth by applying 2,3,5-triiodobenzonic acid (TIBA), an auxin transport inhibitor. TIBA inhibited TEs formation in a *Zinnia elegans* cell culture system (Yoshida et al., 2005). Similarly, TIBA inhibited *de novo* TE formation during *N. benthamiana* grafting (Figure 1, E and F). This inhibition was also reflected in scion growth after 14 DAG (Figure 1, G and H). These results suggested that scion growth relies on *de novo* TE formation and xylem reconnection.

**Figure 1.**
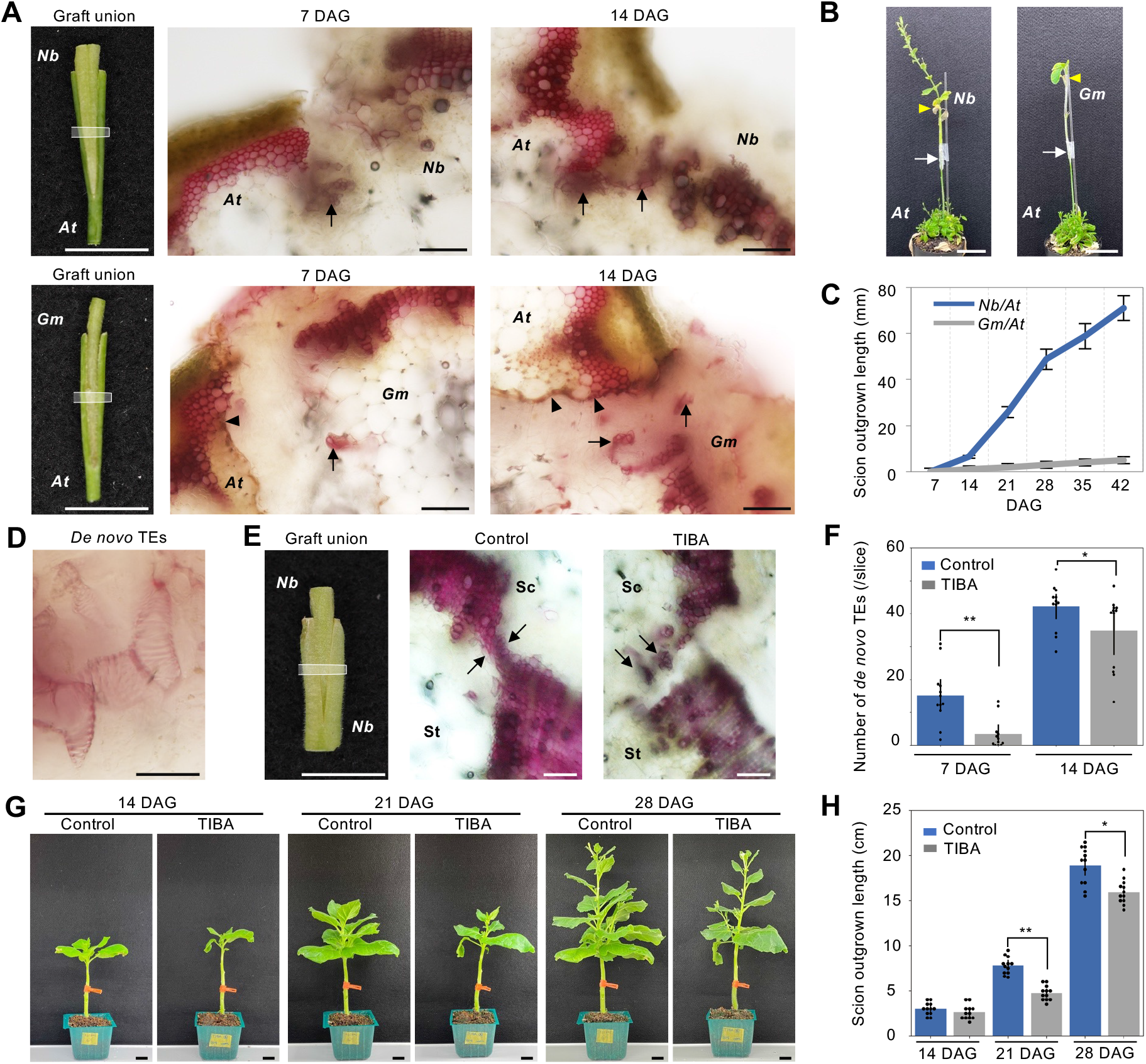
Formation of *de novo* tracheary elements (TEs) at the graft junction. A, Graft union in the *Nb*/*At* and *Gm*/*At* interfamily grafting. Cross sections were made at the middle area of the grafted part (white frame on left) and observed after phloroglucinol-HCl staining to color xylem cells in red, 7 and 14 days after grafting (DAG). Arrows indicate newly generated *de novo* xylem cells. Black arrowheads indicate necrotic layers. Scale bars, 1 cm in the left panels and 100 μm in the middle and right panels. B, Photographs of the interfamily grafts at 42 DAG. Scale bars, 5 cm. White arrows indicate grafted positions. Yellow arrowheads indicate the original apical positions of the scions. C, Length of scion outgrown parts in the interfamily grafts. Elongation of the stem of 36 plants for each graft combination was measured from 7-42 DAG. D, *De novo* TEs formed in the *Nb* calli of the *Nb*/*At* grafts. Scale bar, 20 μm. E, Graft union in self-grafting of *N. benthamiana* in 0.1% DMSO with or without 100 μM of 2,3,5-triiodobenzoic acid (TIBA), an auxin transport inhibitor. Cross sections were made and observed as done for A. Sc, scion; St, stock. Scale bars, 1 cm in the left panel and 200 μm in the middle and right panels. F, The number of *de novo* TEs was counted at 7 and 14 DAG for the samples in E. Four to five cross-sectional slices per graft union were made at region represented by white frame in E. Twelve grafts were measured for each sample fraction. G, H, Post-scion growth for the samples in E was observed (G) and measured (H). Controls in E-H indicate 0.1% DMSO. *At*, Arabidopsis; *Nb*, *N. benthamiana. Gm*, *G. max*. Error bars indicate the means ± SE. Asterisks present significant differences determined by Student’s T-test results (\**P* < 0.05; \*\**P* < 0.01).

### Xylem formation genes were expressed during graft establishment

We analyzed the transcription of genes related to xylem formation to investigate the molecular mechanism of xylem formation during interfamily grafting. In previous research on Arabidopsis, *VND1–VND7* were considered master transcription factors that initiate xylem formation processes, such as cell autolysis and cell wall synthesis, through the regulation of downstream genes (Figure 2A) (Kubo et al., 2005; Ohashi-Ito et al., 2010; Endo et al., 2015; Yamaguchi et al., 2011). Before analyzing gene expression patterns, we examined the timing of xylem formation initiation. We conducted *Nb*/*At* grafting to test the establishment of water transport by adding toluidine blue solution to Arabidopsis stock plants and observing dye transport to the scion part by making a section. At 14 DAG, all grafts exhibited xylem bridges, as shown in Figure 1A, and experiments were performed at 1, 2, 3, 7, and 10 DAG, in which 8—10 grafts were examined at each time point. Dye transport was first detected at 3 DAG, 20% (2 out of 10) of grafts. The detection frequency did not change at 7 DAG and increased to 62% (6 out of 8) of grafts at 10 DAG (Figure 2B), suggesting that xylem formation started around 3 DAG and kept developing with time. We examined the gene expression patterns at 0, 1, 3, and 7 DAG in successful *Nb*/*At* and unsuccessful *Gm*/*At* interfamily grafts (Figure 2, C and D) based on this observation. All four *VND7* homologous genes in *N. benthamiana* (*NbVND7s*) were upregulated more than the other *VND* homologs in the *Nb*/*At* grafts, indicating that these genes are important for *de novo* xylem formation at the graft boundary in *Nicotiana* (Figure 2C). In the *Gm/At* grafts, one of the two *VND7* homologous genes in *G. max* was upregulated but the other was not (Figure 2D). To examine the expression patterns of downstream genes of *VND7s* in these grafts, homologs of 19 genes known as *VND7* downstream genes in Arabidopsis (Kubo et al., 2005; Ohashi-Ito et al., 2010; Endo et al., 2015; Yamaguchi et al., 2011) were identified in *N. benthamiana* and *G. max* by phylogenetic analysis. Most genes involved in xylem formation were upregulated in the *Nb*/*At* grafts but downregulated in the *Gm*/*At* grafts (Figure 2, C and D). Thus, the morphological features of TE formation at the graft junction are consistent with the expression patterns of xylem formation-associated genes.

**Figure 2.**
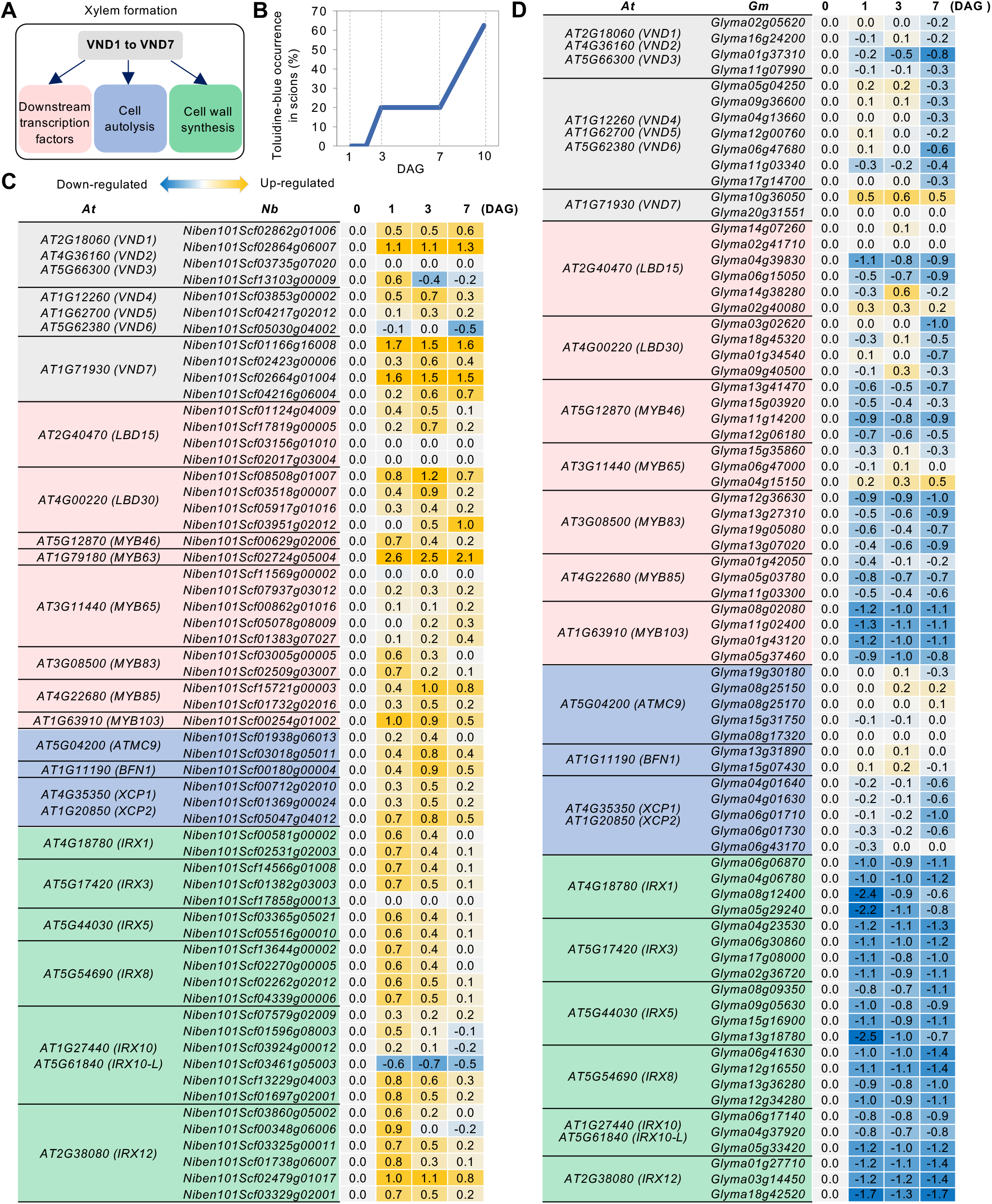
Expression profiles of VNDs-initiated xylem formation genes. A, Proposed VNDs-initiated processes in xylem formation. *VND1–VND7* are master transcription factors that regulate various processes of xylem formation, according to Arabidopsis research (Kubo et al., 2005; Ohashi-Ito et al., 2010; Yamaguchi et al., 2011; Endo et al., 2015). B, Measurement of the vascular tissue reconnection in the *Nb*/*At* interfamily grafts. The frequency of toluidine-blue detection in the scion was measured at 1, 3, 5, 7, and 10 DAG. Eight to ten grafts were examined for each time point. C, D, Expression profiles of xylem formation-related genes in *N. benthamiana* and *G. max*. The transcription levels of genes in *Nb/At* or *Gm*/*At* interfamily grafting were estimated by RNA-seq. Colors in the heat map represent the fold changes (log10) in the transcription level at 1, 3, and 7 DAG relative to the intact stem (0 DAG). Xylem formation-related genes in Arabidopsis were listed from previous study (Yamaguchi et al., 2011; Bollhöner et al., 2013; Ohashi-Ito et al., 2018). Corresponding genes in *N. benthamiana* and *G. max* were orthologous genes of Arabidopsis estimated by the tblastn program. *At*, Arabidopsis; *Nb*, *N. benthamiana; Gm*, *G. max*.

### Network analysis revealed a core module for xylem differentiation during grafting

To investigate the genes involved in xylem formation during grafting, we performed gene network analysis. We first performed Weighted Gene Correlation Network Analysis (WGCNA) using transcriptome data from 3 replicates of intact *N. benthamiana* stems and *Nb*/*At* grafting sites at 2 hours after grafting, 1, 3, 5, and 7 DAG (Supplemental Figure S2). The iDEP 0.96 interface (http://bioinformatics.sdstate.edu/idep96/) was used to draw co-expression networks for 3,000 most variable genes (Supplemental Figure 2A). A “soft threshold” of “9” and a “minimum module size” of “100” generated 6 well-defined modules consisting of 2,818 genes (Supplemental Figure S2B–D). These 6 modules contained 11 genes related to canal formation as shown in Figure 2C (Supplemental Table S1). From the *N. benthamiana VNDs* were included in modules 2 and 6, we extracted 313 genes that were linked neighboring to the two respective *VND7* within those modules (Supplemental Table S2). For those genes, gene ontology analysis was performed using Arabidopsis homolog annotation. We detected transcription factors in the molecular function category, cell wall-related genes in the cellular component category, and water deprivation and wound response genes in the biological process category (Supplemental Table S3). However, this analysis did not resolve the causal relationship of gene expression, so it is not possible to distinguish whether these co-expressed genes are regulated by *VND7* or affect the expression of *VND7*.

We next performed a Bayesian network analysis, which can reveal the causal relationship between gene expression, using the SiGN-BN NNSR program (Tamada et al., 2010; 2011) with 36 *N. benthamiana* grafting transcriptome data (Notaguchi et al., 2020). All data from the whole-genome transcriptome were used as initial inputs to draw the gene network (Figure 3A). In the network, 2,038 *N. benthamiana* genes were associated with four *NbVND7s* within the sixth level (a direct connection is the first level, a connection bypassing one other gene is the second, and so on) (marked in yellow in Figure 3A) including *LBD15*, *LBD30*, *ATMC9*, *Myb83 VND2*, *XCP1*, *XCP2* and various unidentified genes (Supplemental Table S4). To further analyze the gene regulatory network of xylem differentiation, we attempted to apply the Bayesian network using the SiGN-BN HC+Bootstrap program (Tamada et al., 2010; 2011) that can describe upstream/downstream relation among genes. Since the upper limit of input in this program is about 1,000 genes, we narrowed down the genes to focus on through gene ontology (GO) analysis (see detail in Supplemental Text 1 and Supplemental Table S5). The Bayesian network for four *NbVND7s* and 846 *N. benthamiana* genes categorized into four GO groups were drawn (Figure 3B, Supplemental Table S6). By narrowing down the genes associated with *de novo* xylem formation during grafting, we highlighted a core structure from *NbVND7s* to genes for xylem formation among the entire network linking to *NbVND7s*. We compared the 846 *N. benthamiana* genes with the 189 *N. benthamiana* graft-induced genes at 1—3 DAG that were previously identified since xylem formation was initiated after 3 DAG (Figure 2B) (Notaguchi et al., 2020). As nine genes overlapped in the comparison (Supplemental Table S7), we extracted the nodes and edges starting from *NbVND7s* and ending with the nine genes as a core network that potentially links to xylem formation (Figure 3C, Supplemental Table S8). In the center of the core network, the overlapping gene *Niben101Scf01015g01002*, a homolog of *pathogenesis-related 4* (Camargo-Ramírez et al., 2018) and named *NbPR-4* is connected by several *NbVND7* downstream genes. The other eight overlapping genes are at the bottom left of the network and have strong interactions among themselves. By ranking the top 100 edges based on relationship strength, two distinct clusters were identified, including all nine overlapping genes (Figure 3D). One cluster consisting of eight overlapping and eight additional genes included genes expected to be directly involved in tracheary element differentiation, such as all four Arabidopsis *xylem cysteine peptidase 1* and *xylem cysteine peptidase 2* (*XCP1* and *XCP2*) orthologs, named *NbXCP1–4*, a *tracheary element differentiation-related 6* (*TED6*)ortholog, and a *glycosyl hydrolase family 10 protein* (*ATXYN1*) ortholog (Avci et al., 2008; Endo et al., 2009; 2019), and the other cluster connecting to *NbPR-4* consists of an additional six genes, whose roles in xylem formation are unknown (listed in Figure 3E and see also Supplemental Text 2).

**Figure 3.**
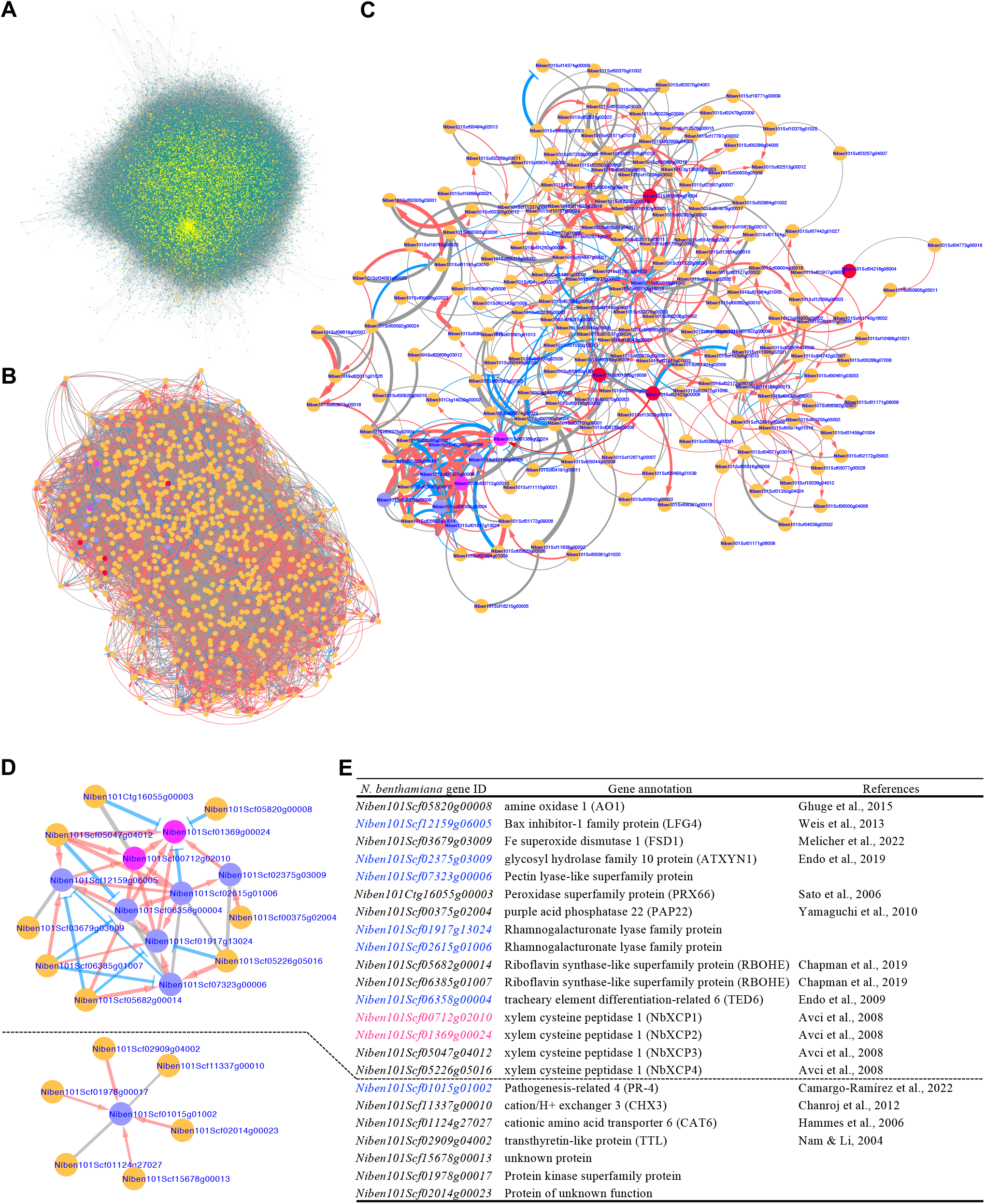
Bayesian network analysis of gene clusters under the control of *NbVND7* transcription factors. A, Bayesian network of all genes in *N. benthamiana* estimated using the SiGN-BN NNSR program. Yellow dots indicate the candidate genes of *NbVND7* downstream to the fifth level. B, Bayesian network of *NbVND7* and genes belonging to the four GOs (Integral component of membrane, Plasma membrane, Extracellular region, and Membrane) included in the downstream genes of *NbVND7* estimated using the SiGN-BN HC+Bootstrap program. C, Network diagram with only the edges and nodes containing graft-related genes extracted. Red and magenta circles indicate homolog genes for *VND7* and *XCP1* in *N. benthamiana*, respectively. Purple circles indicate other graft-related genes. D, Network diagram drawn with the two most significant edges. E, Gene list for network analysis. *Nb*, *Nicotiana benthamiana*. Red arrows indicate gene up-regulation. Blue T-lines indicate gene down-regulation. Gray lines indicate uncertain gene regulation. Line thickness indicates significance in gene regulation (B—D).

### *NbXCP* genes are expressed to form *de novo* TE during grafting

To validate the described gene regulatory network during grafting, as an example, we tested the role of *NbXCP* homologs that were supposed to degrade cellular contents through protease activity to accomplish mature TE formation (Avci et al., 2008) on xylem formation at the graft junction. *NbXCP1* and *NbXCP2* were most similar (99% similarity in coding sequence region), followed by *NbXCP3* (Figure 4A and Supplemental Figure S3A). *NbXCP4* has a large C-terminal deletion and is likely to be a pseudogene that does not functin as a protein (Supplemental Figure S3A). *NbXCP1–3* were upregulated after interfamily grafting, but *NbXCP3* expression was lower (Figure 4B). *NbXCP3* positively affected the expression of *NbXCP1* and *NbXCP2* (Figure 3D), suggesting that the control of *NbXCP3* expression may be different from those of *NbXCP1* and *NbXCP2*. Therefore, we focused on *NbXCP1* and *NbXCP2*. We conducted a heterogeneous yeast one-hybrid assay using the *NbXCP1* promoter (as a representative) and Arabidopsis VNDs to verify the control of *NbXCP* gene expression by *VND* homolog transcription factors. Only VND7 could bind to the *NbXCP1* promoter (Supplemental Figure S4, Figure 4C). The target sequence of VND7 (Tamura et al., 2019) was identified in the *NbXCP1* promoter (Figure 4C). An equivalent sequence was present in the *NbXCP2* promoter, although it contained a mismatch of one terminal nucleotide.

**Figure 4.**
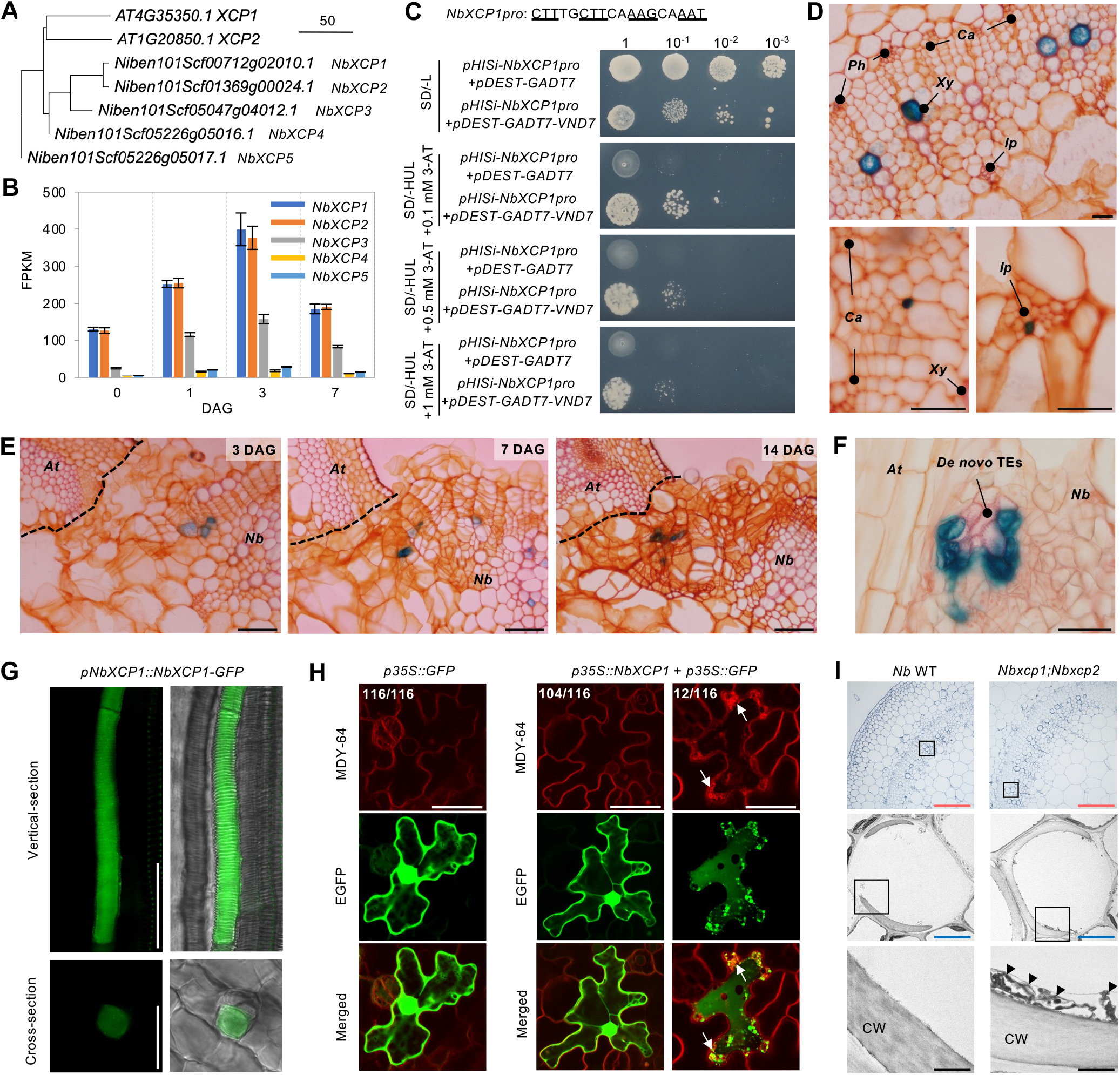
Expression of *NbXCPs* during the graft union formation. A, Phylogenetic tree for XCP1 homologs from Arabidopsis and *N. benthamiana*. B, Expression of *NbXCPs* during graft union formation. The transcription levels of genes in *Nb*/*At* interfamily grafting were estimated by RNA-seq at 0, 1, 3, and 7 DAG. 0 DAG was an intact plant without grafting. Error bars indicate the means ± SE. C, Yeast one-hybrid assay for *VND7* binding to *NbXCP1* promoter. The sequence for *VND7* binding in the *NbXCP1* putative promoter area are shown. Base pairs with underlines indicate a conserved structure ideal for binding by *VND7* protein. Dilution analysis of the yeast transformants harboring *pHISi-NbXCP1 pro* and empty *pDEST-GADT7* vectors or *pHISi-NbXCP1 pro* and *pDEST-GADT7-VND7* vectors on Synthetic Defined (SD) medium. –L: SD medium lacking Leu; –HUL: SD medium lacking His, Ura, and Leu; 3-AT: 3-amino-1,2,4-triazole for reducing *HIS* leaky expression. D, *NbXCP1* promoter directs GUS expression at 4-week-old intact plant stems. Xylem cells were stained in pink and other tissue cells in orange by Safranin-O. Scale bars, 50 μm. *Ph*: phloem; *Ca*: cambium; *Xy*, xylem; *Ip*: internal phloem. E, *NbXCP1* promoter directs GUS expression during *Nb*/*At* interfamily graft union formation at 3, 7, and 14 DAG. The dotted lines indicate the contact surface of *N. benthamiana* and Arabidopsis. Scale bars, 100 μm. F, GUS expression in *de novo* TEs. The putative promoters of *NbXCP1* directed GUS expression at 14 DAG. Scale bars, 50 μm. G, GFP expression in the xylem vessel. The putative promoters of *NbXCP1* directed NbXCP1-GFP fusion proteins (*pNbXCP1::NbXCP1-GFP*) expression at 4-week-old plant stems. Scale bars, 100 μm. H, Ectopic expression of *NbXCP1*. *Nb* epidermal cells were particle-bombarded with the plasmid vectors and observed after one day. MDY-64 stained vacuole membrane. Arrows indicate the anomalous membranes within epidermal cells. “The number of cells with GFP localization pattens represented in each image / total number of the observed cells” are indicated in images. Scale bars, 50 μm. I, Transmission electron microscopic (TEM) observations of tracheary elements (TEs). Observation of TEs in the primary xylem of the stems. TEs were observed in 4-week-old plants. Rectangles indicate the magnified area for continuous observation. Arrowheads indicate remnants attaching cell wall (CW). Scale bars: red, 100 μm; blue, 10 μm; black, 1 μm. *At*, Arabidopsis; *Nb*, *N. benthamiana*.

To investigate *NbXCP1* and *NbXCP2* expression patterns in plant tissues, we generated promoter GUS transgenic lines (Supplemental Figure S5). In *pNbXCP1::GUS* and *pNbXCP2::GUS* lines, GUS expression was found in the xylem tissues of seedlings (Supplemental Figure S6) and the xylem and internal phloem of the stems of 4-week-old plants (Figure 4D, Supplemental Figure S7). GUS expression is restricted to TEs rather than to other xylematic cell types. We conducted a GUS expression analysis in *Nb*/*At* interfamily grafting. GUS expression was concentrated in the TEs newly generated in the callus at the graft junction (Figure 4, E and F, Supplemental Figure S7). To estimate protein function, we generated GFP fusion lines to examine the subcellular localization of NbXCP1 and NbXCP2 (Supplemental Figure S5). When NbXCP1-GFP and NbXCP2-GFP fusion proteins were expressed in the stem using their promoters, GFP fluorescence was detected in the cytoplasmic region of TE maturating cells (Figure 4G, Supplemental Figure S7). Except for the xylem cells, we detected the GUS stains and GFP expression in the mature pollen (Supplemental Figure S8), the autolysis of which has been reported in Arabidopsis, *Lycopersicum peruvianum*, *Olea europaea*, *Lolium perenne* (Yamamoto et al., 2003; Pacini et al., 2011). We performed particle bombardment to express GFP alone or NbXCP1 and GFP simultaneously in *N. benthamiana* leaf epidermal cells. For the GFP solely expressed, GFP fluorescence was detected in the cytoplasmic region and nucleus in all 116 cells observed. In contrast, for the NbXCP1 and GFP simultaneously expressed, ~10% (12/116) of cells exhibited a planar GFP fluorescence pattern with strong GFP dots, indicating cytoplasmic degradation in the cells (Figure 4H). Finally, we confirmed the function of *NbXCP1* and *NbXCP2* in TE maturation by analyzing a double knockout mutant of *NbXCP1* and *NbXCP2* generated using the CRISPR/Cas9 system (Supplemental Figure S9). Transmission electron microscopy revealed that the *Nbxcp1;Nbxcp2* mutant showed a defect in cellular digestion in differentiating TEs of the stem compared with the wild-type (WT) (Figure 4I), as observed in the Arabidopsis *xcp1;xcp2* mutant (Avci et al., 2008). Overall, *NbXCP1* and *NbXCP2* play a conserved role in TE formation.

### *De novo* TE formation is essential for graft establishment and post-scion growth

We investigated the function of *NbXCP1* and *NbXCP2* during grafting to determine whether xylem formation is important for the establishment of interfamily grafting. We conducted virus-induced gene silencing experiments as reported previously (Notaguchi et al., 2020). *Cucumber mosaic virus* (*CMV*)vectors containing a 295 bp sequence of *NbXCP1* that is identical to that of *NbXCP2* (*CMV-NbXCP1/2*)or a 292 bp sequence of GFP (*CMV-GFP*) were infected with *N. benthamiana* shoots and grafted onto the Arabidopsis stock. RT-PCR and qRT-PCR analyses demonstrated knockdown of *NbXCP1* and *NbXCP2* (Supplemental Figure S10, Figure 5A). Compared with the non-infected (NI) and *CMV*-*GFP*-infected controls, *CMV*-*NbXCP1/2*-infected *N. benthamiana* scions showed significantly lower survival rates at 14 DAG (Figure 5B). We next examined interfamily grafting using the *Nbxcp1;Nbxcp2* knockout mutant and *NbXCP1* translationally enhanced (*NbXCP1-OX*) lines that maintained endogenous expression patterns by using own promoter (Supplemental Figure S9). Although there was a slight decrease of the fresh weight only in the intact *Nbxcp1;Nbxcp2* mutant seedlings, not in the *NbXCP1*-*OX* lines, the growth retardation was no longer observed in the grown plants (Supplemental Figure S9). The survival rate decreased in the *Nbxcp1;Nbxcp2*/*A*t grafts, while the *NbXCP1-OX*/*At* grafts tended to increase (Figure 5C). The TE formation timing at the graft junction was measured for each graft combination. *De novo* TE formation began 3 DAG. The frequency of grafts that first formed TE at the graft junction was lower in the *Nbxcp1;Nbxcp2* mutant and higher in the *NbXCP1-OX* line than in the WT at 3 and 4 DAG. All grafts formed TEs at 5 DAG or later in all graft combinations (Figure 5D). Consistent with this observation, transport of isotope-labeled phosphorus-32 (^32^P) to the *N. benthamiana* scions was lower in the *Nbxcp1;Nbxcp2* mutant than in the WT and *NbXCP1-OX* plants at 7 DAG (Figure 5E). This finding revealed that *NbXCP1* and *NbXCP2* affect *de novo* TE formation from the early phase of graft union formation and that the pace of *de novo* TE formation at graft junction affects the level of water transport from the rootstock to the scion and scion growth. We continued to grow the grafts and measured scion growth. The scion growth of the *Nbxcp1;Nbxcp2* mutant was lower but the *NbXCP1-OX* scion grew more than the WT *Nb* scion (Figure 5F). The scion growth was also reflected in fruit production. The fresh weight of fruits was decreased in the scion of *Nbxcp1;Nbxcp2* mutant compared to the wild-type *N. benthamiana* scion. In contrast, the fruit weight was increased in *NbXCP1-OX* scion (Figure 5H). However, we did not find the fruit weight has different in intact plants of WT, *Nbxcp1;Nbxcp2* mutant and *NbXCP1*-*OX* line. These findings indicate that the function of *NbXCP1* and *NbXCP2* is important for both scion survival and post-grafting growth.

**Figure 5.**
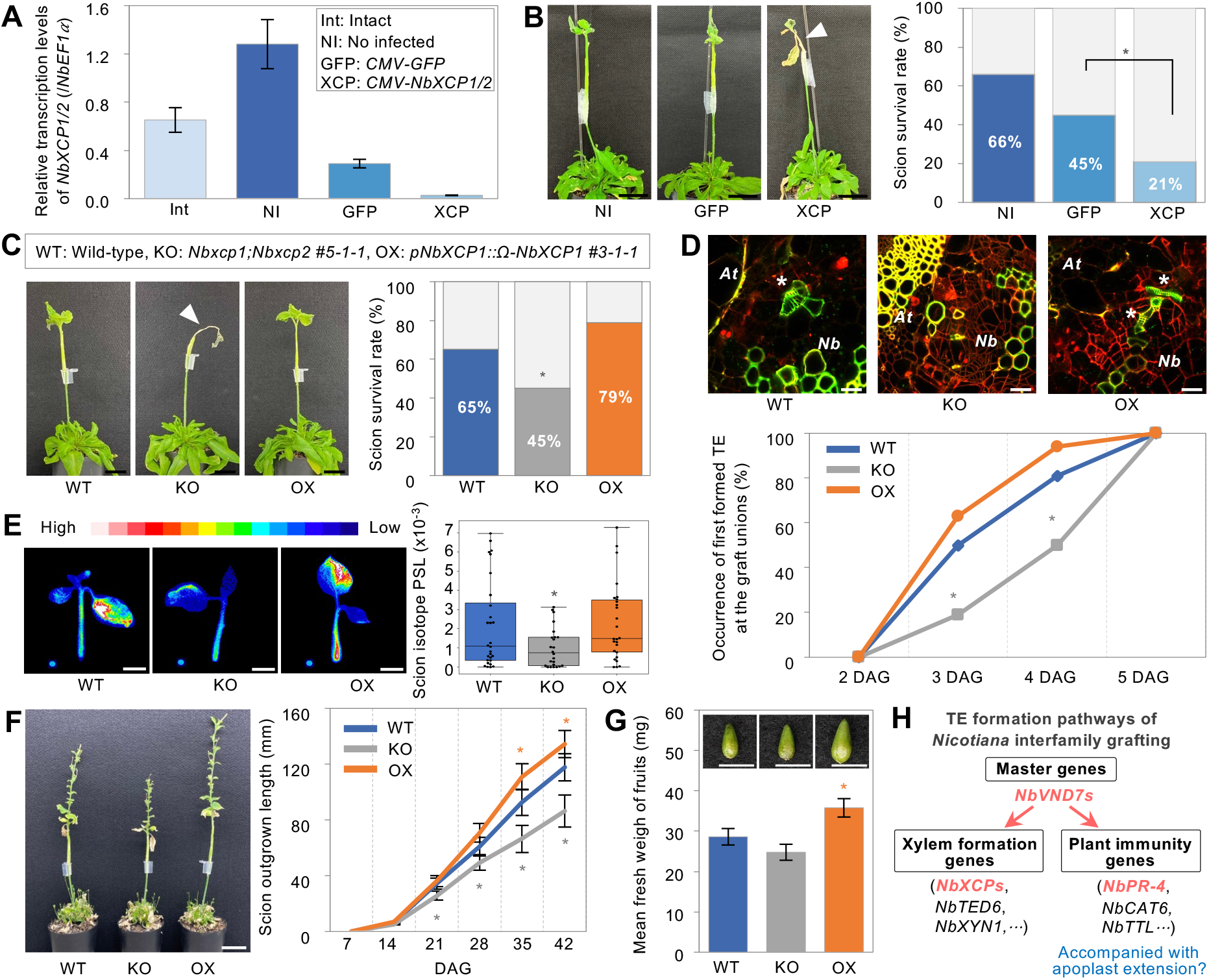
*NbXCP1* and *NbXCP2* promote graft union formation and scion growth. A, Suppression of *NbXCP1* and *NbXCP2* transcripts in interfamily grafting by virus-induced gene silencing (VIGS). *CMV-GFP* and *CMV-NbXCP1/2* indicate infected *Nb* plants with *cucumber mosaic virus (CMV*) containing *GFP* and *NbXCP1/2* fragments, respectively (See also Supplemental Figure S10). The related gene transcription in graft junctions was verified at 3 DAG by qRT-PCR. *NbEF1α* was used as a reference. Error bars indicate the means ± SE (n = 3-4). B, Effect of *NbXCP1* and *NbXCP2 s*uppression on graft establishment. Representative *Nb*/*At* grafts at 14 DAG in VIGS experiment. Arrowhead indicates the dead scion. The survival rates were measured with 46–48 grafts for each at 14 DAG based on whether the *Nb* scion was alive. Asterisks present significant differences determined by the Chi-square test (\**P* < 0.05). Scale bars, 2 cm. *Nb*, *N. benthamiana*; *At*,Arabidopsis. C, Grafts of *Nbxcp1:Nbxcp2* knockout mutant (KO) and *NbXCP1* translationally enhanced (OX) line as a scion on the stems of Arabidopsis at 14 DAG. Survival rates were estimated with 45–48 plants for each at 14 DAG. The arrowhead indicates the dead scion. Scale bars, 2 cm. Asterisks present significant differences determined by the Chi-square test (\**P* < 0.05). D, *De novo* TE differentiation in the calli of graft union at 2 DAG. Propidium iodide (PI) and BF-170 were used for staining dead cells and xylem cells, respectively. The occurrence of *de novo* TEs in calli was estimated with 16 plants for each at 2, 3, 4, and 5 DAG. Asterisks represent significant differences determined by the Chi-square test (\**P* < 0.05). White asterisks indicate *de novo* TEs in calli. Scale bars, 50 μm. E, Radioisotope transport assays in *N. benthamiana* scions. All plantlets incorporated inorganic phosphate (Pi) labeled with ^32^P from the cut stems. The signal amount is shown as a heat map. The gradient from white to blue corresponds to higher to lower signals. Pi transport was quantified with 26–30 scions for each by imaging plates at 7 DAG. Error bars indicate the means ± SE. Asterisks present significant differences determined by Tukey’s HSD test result (\**P* < 0.05). Scale bars, 1 cm. F, Measurement of scion growth of *Nbxcp1;Nbxcp2* knockout mutant and *NbXCP1* translationally enhanced line grafted on Arabidopsis grafts at 42 DAG. Scale bars, 4 cm. Outgrow-part of *Nb* scions with 33–36 plants for each were measured at 7, 14, 21, 28, 35, and 42 DAG and plotted. Error bars indicate the means ± SE. Asterisks present significant differences determined by Tukey’s HSD test result (\**P* < 0.05). G, Measurement of fruit weight of *Nbxcp1;Nbxcp2* knockout mutant and *NbXCP1* translationally enhanced line grafted on Arabidopsis grafts at 49 DAG. Scale bars, 5 mm. The fresh fruits were harvested from 20-24 plants graft combination. The measurement with each single fruit. Error bars indicate mean ± SE (n = 36—69 for each). Asterisks present significant differences determined by Tukey’s HSD test result (\**P* < 0.05). H, A model for *de novo* TE formation in *Nicotiana* interfamily grafting. *NbVND7* transcriptional factors regulate the downstream genes including xylem formation genes and plant immunity genes during the TE differentiation. Bayesian network analysis reveals the candidate genes in graft union formation. Among them, the function of *NbXCP1* and *NbXCP2* on *de novo* TE formation has been verified in this study.

## Discussion

Water uptake from the soil through water-conducting tissues is essential for most land plants. Therefore, in grafting, new xylem formation at the graft junction is necessary for the survival of grafted plants. This study addressed this principle by comparing successful and unsuccessful interfamily grafting, where xylem formation was achieved and completely absent, respectively, and comparing mock and exogenous TIBA treatments to delay the timing of xylem formation. In Arabidopsis hypocotyl grafting, it was shown that TIBA suppresses the cell proliferation of vascular tissue during graft union formation (Matsuoka et al., 2016). Moreover, TIBA and the other auxin transport inhibitors, also blocked the formation of TEs and continuous vascular stands at the interface of the haustorium and the host tissues (Yoshida et al., 2005; Wakatake et al., 2020). The experiments in this study confirmed that xylem formation is essential for the *N. benthamiana* interfamily grafting. The timing of xylem connections after grafting determine the post-grafting growth of scion plants (Figure 1) and it was accelerated by the overexpression of *NbXCP1*, resulting in the enhancement of scion growth (Figure 5).

Transcriptomic analysis demonstrated that previously known xylem-associated genes exhibited consistent expression patterns (Figure 2). Previous transcriptome analyses on graft junctions also identified gene expression controls of xylem-associated genes (Kurotani et al., 2022; Melnyk et al., 2015; 2018; Thomas et al., 2022). In this study, we estimated a gene network related to xylem formation, whose accuracy was evaluated by analyzing a significant gene, *NbXCP* (Figures 4 and 5), and described the features of the gene modules (Figure 3). In previous studies, two studies on gene network analysis for plant grafting have been reported. A co-expression network was used to identify genes involved in graft formation (Xie et al., 2019). A Bayesian network has been used to detect gene regulatory networks especially involved in upstream event of grafting and successfully identified *SlWOX4* as a key transcription factor to trigger downstream gene expressions (Thomas et al., 2022). In this network analysis, DEG and GO analyses were first conducted to narrow down the target genes for network estimation in order to match the limit of the number of genes that can be used as the input of the network analysis method used. This limitation in gene numbers for input required cutting out a part of gene population. To overcome this number limitation, the SiGN-BN HC+Bootstrap program (Tamada et al., 2010; 2011) was applied to estimate Bayesian networks on a genome-wide scale. An entire network was successfully obtained where expression datasets of all genes of *N. benthamiana* were used as an initial input. However, since we could not solve the causal relationship on this scale, we reduced the number of genes to about 1,000 by capturing our interested portion out of entire gene network, genes associated xylem formation in this study. To eliminate artificial bias as much as possible, the *P* value and population size were used as criterions in this study, rather than the contents of GOs, resulting in identification of an unexpected module (Figure 3).

In this study, we proposed a model (Figure 5H) for TE formation pathways of *Nicotiana* interfamily grafting. There are two reliable relationship gene modules under the control of *NbVND7s*, the master transcriptional element genes; a module involved in xylem formation, including *NbXCPs* and a module involved in plant immunity, centering *NbPR-4*. In the module of xylem formation, we have revealed the orthologs of *tracheary element differentiation-related6* (*TED6*) and *endo-β-1,4-xylanase* (*XYN1*). *TED6* was identified as secondary cell wall (SCW)-related membrane proteins, participating the formation of tracheary element. An interaction was found between the TED6 and a member of the *SCW*–*Cellulose Synthase* (*CesA*) complex (Endo et al., 2009; Rejab et al., 2015). *XYN1* was verified expression on the predominantly in xylem tissues and encodes xylanase. The xylem transport patterns were affected in *AtXYN1* modified lines (Zeilinger et al., 1996; Suzuki et al., 2002; Endo et al., 2019). In the module of xylem formation, genes associated with the generation and scavenging of reactive oxygen species (ROS) were found (Figure 3, Supplemental Table S7). These genes are considered to be directly involved in the xylem formation process. In Arabidopsis, secondary cell wall formation consisting of cellulose, hemicellulose, and lignin has been observed during xylem development. It has been shown that lignification proceeds by polymerization of monolignols, in which peroxidase uses hydrogen peroxide as a substrate (Boerjan et al., 2003; Fagerstedt et al., 2010; Tobimatsu et al., 2019). Thus, this module can be highly associated with xylem cell differentiation.

The module involving *NbPR-4* contains the orthologs of *cationic amino acid transporter 6* (*CAT6*)and *transthyretin-like protein* (*TTL)*. In Arabidopsis, *AtCAT6* was nematode-induced in roots during the infestation, mediating transportation of amino acids to prevent plant injury (Hammes et al., 2006). Research also reported that *CAT6* and *CAT7* involve in oxidative stress response in cassava plants (Wang et al., 2021). The TTL protein was found as scaffold proteins for a potential substrate of *Brassinosteroid-insensitive 1* (BRI1) in Arabidopsis, promoting brassinosteroid responses (Nam et al., 2004; Amorim-Silva et al., 2019). Brassinosteroids play an important role in inhibiting pathogen infection, in which mediating growth directly antagonizes innate immune signaling (Albrecht et al., 2012). However, three of the four genes positively regulating *NbPR-4* encode the uncharacterized protein kinase or the proteins of unknown function. Further investigation may provide us new insights on the components involved in the relation between xylem formation and immune system in plants.

These findings may imply that there is an inevitable relationship between plant immunity and xylem formation at the graft boundary because it is reasonable that plants react after graft wounding to avoid pathogen infection. In fact, the tyloses from xylem parenchyma cells have been shown to give resistance against the pathogens (Grimault et al., 1994; Rahman et al., 1999; Clérivet et al., 2000; Fradin and Thomma, 2006). Recent studies on pathogenesis in Arabidopsis reported that *XCP* genes may assist plant immune-related gene expression and enhance resistance against xylem vessel pathogen (Zhang et al., 2014; Chen et al., 2021). Although XCP proteins did not directly act on plant pathogens (Zhang et al., 2014; Pérez-López et al., 2021), XCP1 activated the systemic immunity by proteolyzing Pathogenesis Related Protein 1 (Chen et al., 2021). Thus, xylem formation, an enlargement of the apoplastic region inside of the plant body, may enhance the plant immune system since it gains potential risks for the exposure to endophytes and pathogenic bacteria transmitted through the xylem tissue. However, further elucidation is required for this hypothesis. The spatiotemporal gene expression patters of potential immune-related genes will be important to understand the mechanism.

Thus, the gene network described in this study may include a set of cellular aspects of xylem formation and plant immune mechanism during grafting. In the future, it would be interesting to investigate newly identified genes during grafting and/or xylem formation in the development of other organs. In addition, the similarity of the network of TE formation between interfamily and intrafamily grafting using accumulated transcriptome data from other species is interesting. Moreover, network analysis of other biological processes during grafting would help to investigate the molecular events of grafting and identify crucial gene modules. This strategy will enhance our understanding of how plants achieve tissue reunion at graft wound sites.

## Materials and Methods

### Plant materials and growth conditions

Seeds of *N. benthamiana* and Arabidopsis were sterilized with 5% (w/v) sodium hypochlorite (NaClO) solution for 5 min, washed six times with sterile water, and incubated at 4 °C in the dark for three days. Seeds of *N. benthamiana* were sown on half-strength Murashige and Skoog medium supplemented with 0.5% (w/v) sucrose and 1% (w/v) agar. The pH was adjusted to 5.8 with 1 M KOH. Seedlings were cultured on medium for seven days in growth chambers and transplanted into the soil in a growth room. Arabidopsis seeds (ecotype Col-0) were directly surface-sown on the soil. Seedlings of *N. benthamiana* and Arabidopsis were grown at 27 °C and 23 °C with 70% and 30% relative humidity, respectively, and 100 μmol m^−2^ s^−1^ continuous illumination. *G. max* seeds were directly sown in the soil in a 23 °C growth room. Plants in the growth room were watered thrice per week.

### Interfamily grafting

Wedge grafting was performed on the stems of 3-week-old *G. max*, 4-week-old *N. benthamiana*, and 5-week-old Arabidopsis plants, as described previously (Notaguchi et al., 2020). *N. benthamiana* and *G. max* stems were cut and trimmed to a V-shape. The inflorescence stems of Arabidopsis were cut and split from the stem middle at the graft site. The V-shaped stem was inserted into a middle-split stem and fixed using wrapping film (Bemis Parafilm M) or clip to form a graft union. The scions were covered with water-sprayed plastic bags and grown in an incubator at 27 °C under continuous light (30 μmol m^−2^ s^−1^) for 10 d. Afterward, plastic bags were removed from the scions, the grafts were transferred to a 23 °C growth room with 30% relative humidity, and 100 μmol m^−2^ s^−1^ continuous illumination. *CMV*-infected grafts were grown in a 23 °C incubator for 10 days. The other experimental conditions were the same as those mentioned above. The scion outgrowth length was measured from the scion top node to the bottom node of the scion outgrow part (Figure 1, B and C) and recorded every week from 7 DAG (days after grafting) to the next several weeks.

### Construction of plasmid vectors

Genomic DNA was extracted from plant tissues using the DNeasy Plant Mini Kit (Qiagen, Hilden, Germany). Target DNA was amplified by PCR using TaKaRa Ex Taq DNA Polymerase (TaKaRa Bio, Tokyo, Japan). DNA segments were purified using MonoFas DNA Purification Kit I (ANIMOS, Saitama, Japan). Total RNA was isolated from plant tissues using the RNeasy Plant Mini Kit (Qiagen). cDNA was synthesized using SuperScript III First-Strand Synthesis SuperMix (Thermo Fisher Scientific, Waltham, USA). DNA fragments were inserted into plasmid vectors using NEBuilder HiFi DNA Assembly Cloning Kit (New England Biolabs, Massachusetts, USA). The primers used in this study are listed in the Supplemental Table S9. For the GUS-fusion construction (Supplemental Figure S5), the putative promoter sequence amplified from the *N. benthamiana* genome was fused upstream of the *GUS* (*β*-glucuronidase) reporter gene in *pENTR* vector (pENTR/D-TOPO Cloning Kit, Thermo Fisher Scientific, Waltham, USA) and transferred into the binary vector *pGWB1* (Nakagawa et al., 2007) based on LR recombination reaction. For the GFP-fusion construction (Supplemental Figure S5), the promoter sequences and the coding sequences (CDS) were fused upstream of *GFP* (green fluorescence protein) reporter gene in *pENTR* vector and transferred into the binary vector *pGWB1* (Nakagawa et al., 2007). For the *NbXCP1* translationally enhanced (*NbXCP1-OX*) construction (Supplemental Figure S5), *pNbXCP1::Ω-NbXCP1*, an artificially synthesized *Ω* (omega) fragment was inserted between the promoter sequence and CDS. To the CRISPR/Cas9 vector, fragments of *NbXCP1/2* were as single guide RNA (sgRNA) added to *pKI1.1R* (Tsutsui et al., 2017), according to the above-described methods.

### Production of transgenic plants

The relevant plasmid vectors (*pGWB1* and *pKI1.1R*) were transferred into *N. benthamiana* by the Agrobacterium-mediated transformation method (Krügel et al., 2002; Weigel et al., 2006) for generating transformants. Transgenic plants of GUS-fusion lines, GFP-fusion lines and *NbXCP1* translationally enhanced (*NbXCP1-OX*) lines were screened by antibiotic resistance in seedlings. CRISPR/Cas9-induced mutants were confirmed by sequencing (Dehairs et al., 2016). Homozygous lines of *NbXCP1-OX* lines and mutants were harvested in T2 or T3 generation, which were used in this study.

### Chemical staining and microscopy

Phloroglucinol-HCl stain solution were composed of one volume of 1% (w/v) phloroglucinol dissolved in 70% ethanol and five volumes of 5 M HCl. The toluidine blue stain solution was 0.01% (w/v) dissolved in 0.1 M NaOAC at pH 4. The safranin-O stain solution was 2.5% (w/v) dissolved in 99% ethanol. Fresh tissue sections were directly cut by hand with a blade. Resin-embedded tissue sections were prepared with Technovit 7100 Kits (Kulzer, Wehrheim, Germany) and cut with a rotary microtome (RX-860, Yamato Kohki, Saitama, Japan). Tissue sections were soaked in stain solution for 5 min and observed using a microscope (BX53, Olympus, Tokyo, Japan) equipped with a digital camera (DP73, Olympus) for high-magnification images.

The GUS staining solution consisted of 10 mM PBS at pH 7, 0.1 % Triton X-100, 10 mM EDTA, 0.5 mg/mL X-Gluc, 0.5 mM potassium ferricyanide, and 0.5 mM potassium ferrocyanide. Plant tissues were placed in a 90% acetone solution for 5 min, rinsed in 1 mM PBS, transferred to GUS stain solution, and incubated at 37 °C for 6–12 hours. Plant tissues were soaked in 70% ethanol to stop GUS staining and microscopically observed or prepared in resin-embedded sections. GUS-stained seedlings were observed using a zoom stereomicroscope (SZX10, Olympus) equipped with a digital camera (DP22, Olympus) for photography.

For cellular GFP observation, tissue sections were treated with ClearSee (FUJIFILM Wako Chemicals, Miyazaki, Japan) solution using a fixative solution. A confocal laser scanning microscope (FV3000, Olympus; LSM5 Pascal, Zeiss, Jena, Germany) was set at 488 nm excitation wavelength. For tracheary element observation, the tissue sections were soaked in advance with a mixed solution composed of 20 μg/mL BF-170 (FUJIFILM Wako Chemicals) solution and 2 μg/mL propidium iodide (P1304MP, Thermo Fisher Scientific) solution. The excitation wavelength of the confocal laser scanning microscope (FV3000, Olympus; LSM5 Pascal, Zeiss) was set to 488 nm for BF-170 and 543 nm for propidium iodide. For transmission electron microscopy (TEM) observations, the plant samples were prepared in pieces and sent to Tokai Electron Microscopy, Inc. for photographs.

### Gene expression analysis

RNA-seq data collected in a previous study (Notaguchi et al., 2020) were used in this study. The RNA-Seq data are available from the DNA Data Bank of Japan (DDBJ; http://www.ddbj.nig.ac.jp/) under accession number DRA009936. qRT-PCR reactions were performed using the KAPA SYBR FAST qPCR Master Mix (2X) Kit (KAPA Biosystems, Wilmington, USA) on the QuantStudio 3 Real-Time PCR System (Thermo Fisher Scientific) with standard protocol. The primers used for PCR amplification are listed in the Supplemental Table S6.

### Network analysis

Whole genome Bayesian network analysis in the normalized datasets of RNA-seq experiment in 24 *Nb*/*At* interfamily grafting, nine *Nb*/*Nb* self-grafting, and three intact *N. benthamiana* stems were performed using a Bayesian network estimation program, SiGN-BN (http://sign.hgc.jp/signbn/index.html) implemented on the supercomputer system at the Human Genome Center of the University of Tokyo (https://supcom.hgc.jp/english) (Tamada et al., 2010; 2011). The estimated gene network was analyzed using the gene network analysis software Cytoscape. GO enrichment analysis was performed with DAVID (https://david.ncifcrf.gov) using Arabidopsis gene IDs.

### Yeast one-hybrid assay

A 1238 bp upstream sequence from the first nucleotide of the start codon was selected for the *NbXCP1* putative promoter. This sequence was ligated to the *pHISi* vector. The CDS sequences of *VND1–VND7* were also ligated to the *pDEST-GADT7* vector. The two plasmid vectors were co-transformed into yeast cells. Transformants were grown on SD (synthetic defined) medium lacking leucine or histidine as reported previously (Hossain et al., 2010; Bass et al., 2016).

### Virus-induced gene-silencing (VIGS) experiments

The plasmid vectors *CY1*, *CMV-Al*, and *CY3* contained tripartite components of *cucumber mosaic virus* (*CMV*) genomic RNA (RNA1, RNA2, and RNA3, respectively). *CMV-Al-NbXCP1/2* and *CMV-Al-GFP* (Supplemental Figure S10A) are genetically modified vectors for generating mutated RNA2. Plasmid vectors were linearized to generate templates for viral RNA *in vitro* transcription (T7 RNA Polymerase System, Takara Bio, Shiga, Japan). RNA1 to RNA3 were equally mixed to infect 3-week-old *N. benthamiana* plants using the mechanical inoculation method (Mochizuki et al., 2009). One week later, newly grown leaves with a mosaic phenotype were used to detect viral infection through RT-PCR (Tanase et al., 2019). The primers used for PCR amplification are listed in the Supplemental Table S9. Virus-infected leaves were ground homogeneously and used for sub-inoculation on 3-week-old plants following the above experiment steps. These infected plants were then used for interfamily grafting after one week of growth.

### Measurement of toluidine blue transport

Interfamily grafts of *Nb*/*At* were used to detect toluidine blue appearing on xylem at 1, 2, 3, 7 and 10 DAG. The bottom of Arabidopsis inflorescence stems was cut and soaked into 0.5% (w/v) toluidine blue solution for 12 hours. Cross sections of scion stems were directly cut by hand with a blade and observed using a microscope (BX53, Olympus, Tokyo, Japan) equipped with a digital camera (DP73, Olympus) for high-magnification images.

### Measurement of radioisotope phosphorus-32 (^32^P) transport

Interfamily grafts of *Nb*/*At* were used to detect ^32^P absorption 7 DAG. The bottom of Arabidopsis inflorescence stems was injected with 2 mL of 0.1μM Pi solution (H_3_^32^PO_4_, 10 kBq/mL), followed by plant absorption for 6 h. Plants were transferred to an imaging plate for 2–3 hours of exposure. The radioactivity of ^32^P was detected. Photo-stimulated luminescence (PSL) divided by area (PSL/area) indicates the numerical value of the ^32^P luminescent signal. The final scion radioactivity value was calculated using the following formula: scion (PSL/area) / rootstock (PSL/area).

## Supporting information

Huang_et_al_Supplemental_Material

Huang_et_al_Supplemental_Tables

## Author contributions

CH, KK, and MN conceived the study and designed the experiments. CH performed the grafting experiments and microscopy analysis. CH and RT performed the VIGS experiments. CH and KK analyzed the transcriptome data. KK conducted the network analyses. NM conducted the yeast one-hybrid experiments. RS and KT performed radioisotope experiments. CH, KK, and MN wrote the manuscript.

## Acknowledgments

We thank M. Hattori, M. Matsumoto and F. Tobe for their technical assistance. We are grateful to C. Masuta (Hokkaido University, Japan) for providing CMV vectors. This work was supported by grants from the Japan Society for the Promotion of Science Grants-in-Aid for Scientific Research (20H03273, 21H00368 and 21H05657 to MN and 22K06181 to KK), Japan Science and Technology Agency (JPMJTR194G to MN), and China Scholarship Council (CSC; No. 201908050204 to CH).

## Conflict of interest

We declare no conflict of interest.

## Notes

### Competing Interest Statement

The authors have declared no competing interest.

