## Supplementary material for "*Nicotiana benthamiana XYLEM CYSTEINE PROTEASE* genes facilitate tracheary element formation in interfamily grafting": Huang_et_al_Supplemental_Material

- **Supplemental Text 1 & 2**
- **Supplemental References**
- **Supplemental Figures 1–10**

### Supplementary Text 1

The genes to focus were narrowed down through GO analysis using the Arabidopsis homologs of the 2038 *N. benthamiana* genes associated with four *NbVND7s* within the sixth level in Bayesian network for whole transcripts was performed (Supplemental Table S5). The collected genes were included in GOs that satisfy two conditions; being significantly enriched at $P < 0.05$ and consisting of more than 100 genes with Arabidopsis gene homologs. In the categories of molecular functions, biological processes and cellular components, 4, 1, and 4 corresponding GOs were detected from 28, 69, and 19 GOs, respectively. The four GOs for molecular functions include "protein binding", "ATP binding", "Transcription factor activity, sequence-specific DNA binding" and "DNA binding". Although they are expected to be mainly involved in the transcriptional regulatory network, we did not target them in this study because we focused on genes that are more closely related to execution factors of xylem formation. For the biological processes, GOs related to cell differentiation and stress response were found but their respective populations in the above criteria were not sufficiently large for further analysis. For cellular components, four GO terms, "Integral component of membrane," "Plasma membrane," "Extracellular region," and "Membrane" were detected. We then conducted further analysis on these 864 *N. benthamiana* genes.

In this study, we focused on major populational GOs. However, GOs that were not used in the analysis due to small population size but were significantly enriched are described below. In the category of molecular function, "UDP-glycosyltransferase activity", "transferase activity, transferring glycosyl groups" and "UDP-glucosyltransferase activity" were significantly enriched at $P < 0.01$. These GOs include cellulose synthase genes such as *cellulose synthase-like A3* (*ATCSLA03*), *ATCSLC12*, and *ATCSLD1*, as well as *galacturonosyltransferase 15* (*GAUT15*), *galactinol synthase 1* (*AtGolS1*), and 10 different *UDP-glucosyl transferases*, indicating that genes involved in cell wall biosynthesis are regulated downstream of *VND7*. In the category of biological process, "mitochondrial fission", "pollen germination", "programmed cell death involved in cell development", "protein phosphorylation", "xylem vessel member cell differentiation", "cell differentiation" and "positive regulation of cellular process" were also enriched at $P < 0.01$. These GOs include transcription factors such as *VND7* itself, *VND2*, *ANAC007*, *ANAC038*, *AGAMOUS-like 6* (*AGL6*), *AGL8*, *AGL20*. As a factor in determining cell fate, *ATMYB5 RGA-LIKE 1* (*RGL1*), which is involved in ROS production, and *AUTOPHAGY 6* (*ATG6*) and *RAB GTPase homolog G3B* (*ATRABG3B*), which regulate autophagy, were found. Other genes involved in cell differentiation such as *Ras-related small GTP-binding family protein* (*ATSGP2*), *Rho GTPase-activating protein enhancer 1* (*REN1*), and *FRIGIDA-like protein* (*FRI*), and many protein kinases such as *calcium-dependent protein kinase 29* (*CPK29*), *CPK33*, *cysteine-rich RLK 19* (*CRK19*), *phosphoenolpyruvate carboxylase kinase 1* (*PPCK1*), *Leucine-rich repeat protein kinase family protein* (*PRK2A*), *receptor lectin kinase* (*RLK*), and *wall associated kinase-like 1* (*WAKL1*) are also included. In the category of cellular components, "cell wall", "intracellular membrane-bounded organelle", "Golgi membrane", "lysosome" and "chromosome" were enriched at $P < 0.01$. These GOs include cell wall-related genes such as *amine oxidase 1* (*AO1*), *xyloglucan endotransglucosylase/hydrolase 3* (*XTH3*), *XTH5*, *xyloglucan xylosyltransferase* 5 (*XXT5*), and *glycosyl hydrolase family 10 protein* (*ATXYN1*) in addition to *cellulose synthases* and *UDP-glucosyl transferases* that overlap with the above categories.

### Supplementary Text 2

A cluster consisting of the eight overlapping and eight additional genes (Figure 3D) includes the genes expected to be directly involved in xylem cell differentiation. All four Arabidopsis *xylem cysteine peptidase 1* (*XCP1*) orthologs, *Niben101Scf00712g02010*, *Niben101Scf01369g00024*, *Niben101Scf05047g04012*, and *Niben101Scf05226g05016,* named *NbXCP1-4*, a *tracheary element differentiation-related 6* (*TED6*) ortholog, *Niben101Scf06358g00004*, and an *ATXYN1* (Avci et al., 2008; Endo et al., 2009; 2019) ortholog, *Niben101Scf02375g03009*. In addition, this cluster included other annotated genes. *Niben101Scf12159g06005* is an ortholog of the Arabidopsis *Bax inhibitor-1 family protein* gene (*LFG4*). It has been suggested that LFG4 may function in plant disease resistance through xylan biosynthesis (Weis et al., 2013). *Niben101Scf07323g00006* encodes a pectin lyase-like superfamily protein that is thought to be involved in cell wall organization, lignin biosynthesis, and phloem and xylem histogenesis. *Niben101Scf01917g13024* and *Niben101Scf02615g01006* both encode rhamnogalacturonate-lyase family proteins with pectin-degrading activity. *Niben101Ctg16055g00003* and *Niben101Scf00375g02004* encode a peroxidase superfamily protein (PRX66) and a purple acid phosphatase 22 (PAP22), respectively. These are genes whose expression regulation has previously been suggested to be downstream of VND7 (Yamaguchi et al., 2010; Sato et al., 2006) *Niben101Scf05820g00008* encodes AO1 (Ghuge et al., 2015). This network indicates that the homologs of PRX66 and AO1, which have oxidative activity, suppress *NbXCP1* expression. *Niben101Scf05682g00014* and *Niben101Scf06385g01007* encode riboflavin synthase-like superfamily proteins (RBOHE), suggesting regulation by the promotion of ROS production (Chapman et al., 2019). *Niben101Scf03679g03009* encodes iron superoxide dismutase 1 (FSD1), which promotes ROS scavenging (Melicher et al., 2022). It has been suggested that these genes regulate a complex combination of positive and negative effects.

On the other hand, the other cluster centering *NbPR-4* includes ion transporter genes, *Niben101Scf11337g00010* and *Niben101Scf01124g27027*, encoding a cation/H+ exchanger 3 (CHX3) (Chanroj et al., 2012) and a cationic amino acid transporter 6 (CAT6) (Hammes et al., 2006), respectively. *Niben101Scf02909g04002* encodes a transthyretin-like protein (TTL) (Nam et al., 2004), indicated to be involved in stress response by catalyzing allantoin biosynthesis. Other genes had unknown functions. Thus, the network analyses revealed stress-induced genes downstream of *NbVND7s*, in addition to the previously known TE formation genes. They were supposed to be closely related to each other.

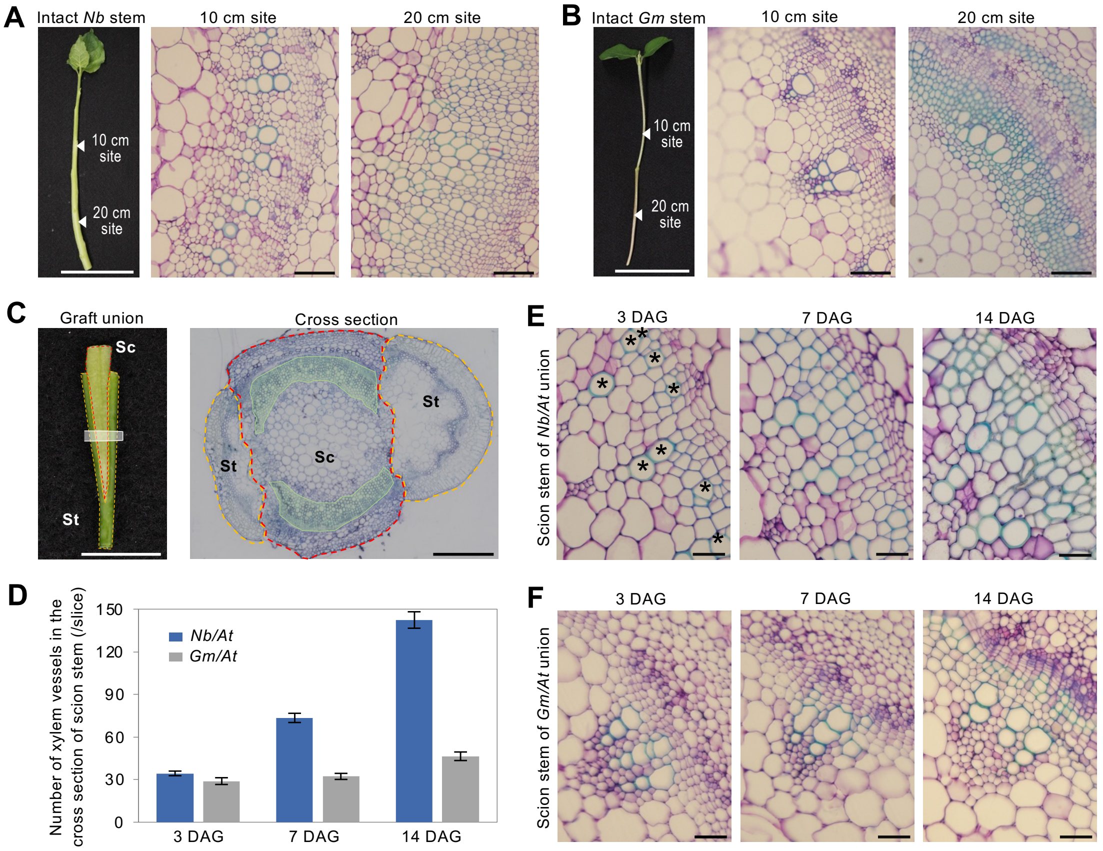


**Supplemental Figure S1** Secondary xylem development in the stems. A, B, Observation of primary xylem and secondary xylem in *N. benthamiana* (*Nb*, A) and *G. max* (*Gm*, B) stems. The section sites were 10 cm and 20 cm from the apex. Toluidine blue was used to stain the xylem blue and other tissues purple. Arrowheads indicate the loci observed by microscopy. Scale bars, white, 10 cm; black, 100 μm. C, Representative photographs of the graft union of *Nb* scion and Arabidopsis (*At*) stock. Red dotted lines indicate scion (Sc) and yellow dotted lines indicate stock (St). The white frame indicates the middle area of graft union for tissue observation. In a represented cross-section, areas colored in green represent the xylem tissues of the scion stem. Scale bars, white, 1 cm; black, 0.5 mm. D, Statistics for the number of xylem vessels in the *Nb*/*At* and *Gm*/*At*. Four graft unions were analyzed for each graft combination at 3, 7 and 14 DAG. Two to three cross sections were sliced from each graft union; the slices were obtained from a middle area of graft union as marked in the white frame in c. By observing cross sections, xylem vessels were counted from regions as marked in green in C. Error bars indicate the mean ± SE. E, F, Xylem development at the graft union of *N. benthamiana* (E) and *G. max* (F) scions in interfamily grafting. The cross sections were stained by toluidine blue. The xylem vessels were marked in asterisks as an example in a photograph of 3 DAG sample in E. Sections were prepared from the area as marked in the white frame in C at 3, 7, and 14 DAG. Scale bars, 50 μm.

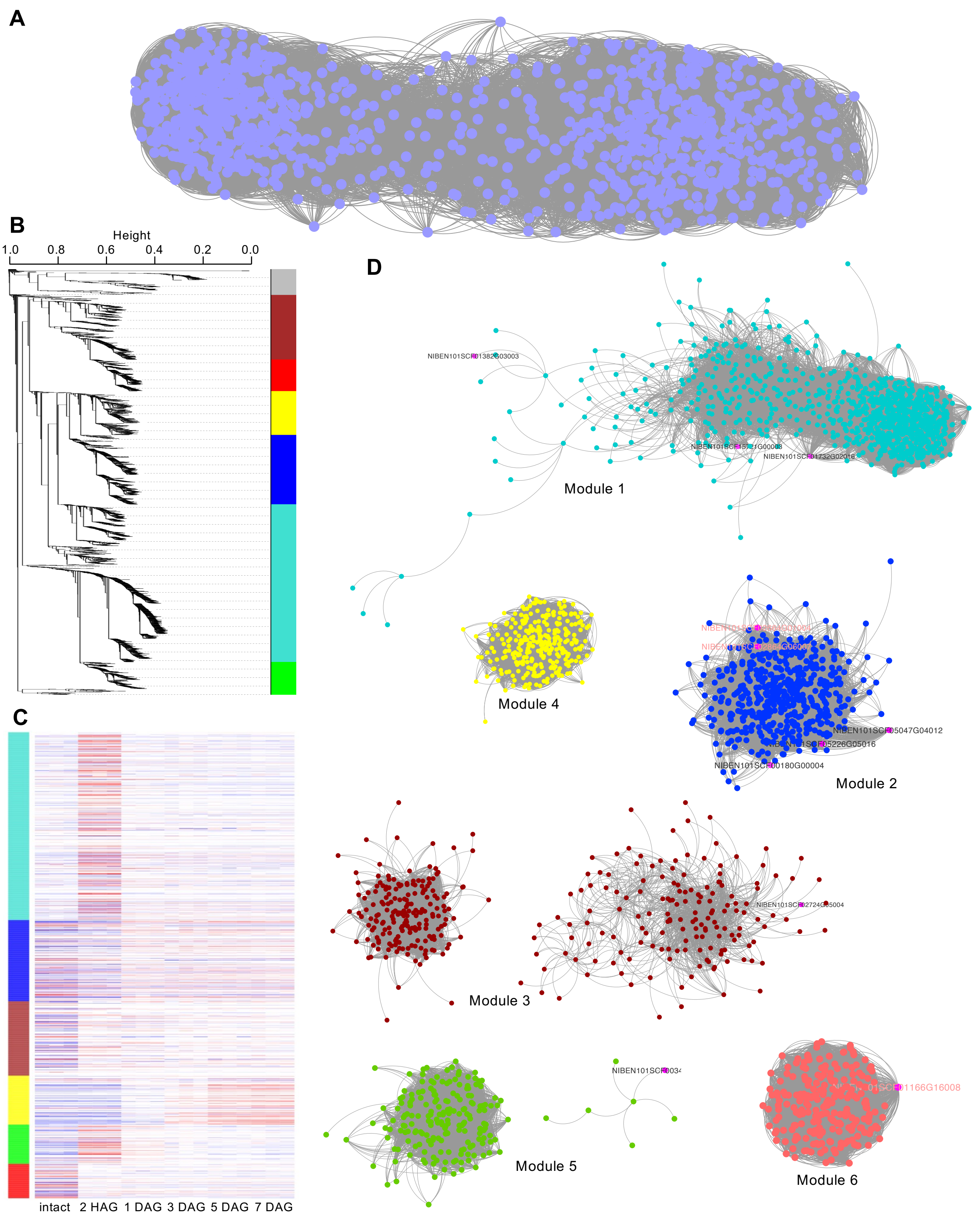


**Supplemental Figure S2** Weighted Gene Correlation Network Analysis in *Nb*/*At* interfamily grafting. A, Weighted Gene Correlation Network Analysis for the most variable 3,000 in *N. benthamiana* estimated using iDEP 0.96. B, Gene dendrogram and modules. The color bars correspond to the modules in C and D. C, Heatmap of identified modules. D, Network diagram drawn with the six modules. Magenta dots indicate the genes related in xylem formation shown in Figure 2. Genes in pink letter indicate *VNDs.*

**A**

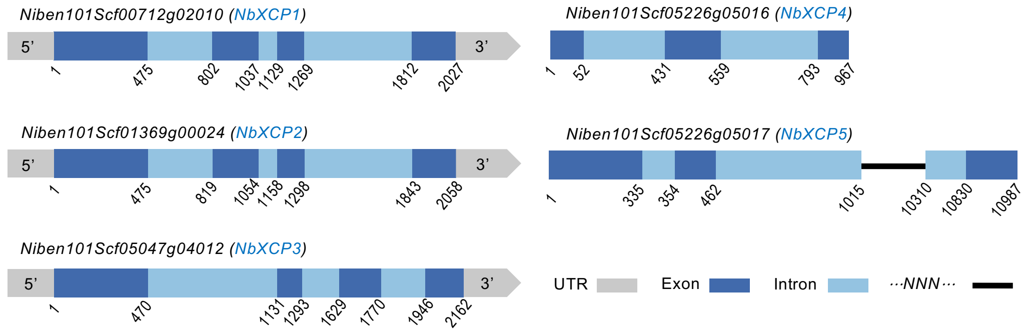


**B**

```
XCP1          MAFSAPSLSKFS-LLVA-ISASALLCCAFARDFSIVGYTPEHLTNTDKLLELFESWMSEH   58
XCP2          MALSS--PSRILCFALALSAASLSLSFASSHDYSIVGYSPEDLESHDKLIELFENWISNF   58
NbXCP1        MAIFS--ISKFS-LLVLFSIF-LLACTTFARDHSIVGYSSDDLTCIDKLINLFEHWMDKH   56
NbXCP2        MAVFS--IFKLS-LLVLFSIS-LLACTSFARDHSIVGYSSDDLTCIDKLINLFEHWMDKH   56
NbXCP3        MASFS--FSKLS-FLALFSILTLVACTTYARDFSIVGYFPDDLNSIENLINLFESWMEKH   57
Conservation  **  :    ::  :           . : ::*.*****  :.*   ::*::*** *:.:.

XCP1          SKAYKSVEEKVHRFEVFRENLMHIDQRNNEINSYWLGLNEFADLTHEEFKGRYLGLAKPQ   118
XCP2          EKAYETVEEKFLRFEVFKDNLKHIDETNKKGKSYWLGLNEFADLSHEEFKKMYLGLKTDI   118
NbXCP1        GKIYKSIEEKLHRFEIFKDNLKHIDETNKIVSNYWLGLNEFADLSHDEFKKIYLGLKVDN   116
NbXCP2        GKIYKSIEEKLHRFEIFKDNLKHIDETNKIVSNYWLGLNEFADLSHDEFKKMYLGLKVDN   116
NbXCP3        SKIYESIEEKLHRFEIFRDNLKHIDERNKMVSNYWLGLNEFADLSHDEFKKMYLGLQIQH   117
Conservation   * *:::***. ***:*::** ***: *:  ..***********:*:***  ****

XCP1          F--SRKRQPSANFRYRDITDLPKSVDWRKKGAVAPVKDQGQCGSCWAFSTVAAVEGINQI   176
XCP2          V-RRDEERSYAEFAYRDVEAVPKSVDWRKKGAVAEVKNQGSCGSCWAFSTVAAVEGINKI   177
NbXCP1        ELLIKKEKSQQEFAYRDFVDLPKSVDWRKKGAVTPVKNQGQCGSCWAFSTVAAVEGINQI   176
NbXCP2        ELLNKREKSQQEFAYRDFVDLPKSVDWRKKGAVTPVKNQGQCGSCWAFSTVAAVEGINQI   176
NbXCP3        E---RSESSPQEFTYRDFVDLPKSIDWRKKGAVTNVKNQGSCGSCWAFSTVAAVEGINQI   174
Conservation        .    :* ***.  :***:********: **:**.*****************:*

XCP1          TTGNLSSLSEQELIDCDTTFNSGCNGGLMDYAFQYIISTGGLHKEDDYPYLMEEGICQEQ   236
XCP2          VTGNLTTLSEQELIDCDTTYNNGCNGGLMDYAFEYIVKNGGLRKEEDYPYSMEEGTCEMQ   237
NbXCP1        KTGNLTSLSEQELIDCDTTYNNGCNGGLMDYAFQFIISNGGLHKEDDYPYLMEEGTCNEK   236
NbXCP2        KTGNLTSLSEQELIDCDTTYNNGCNGGLMDYAFQFIISNGGLHKEDDYPYLMEEGTCDEK   236
NbXCP3        KTGNLTSLSEQELIDCDTKYNSGCNGGLMDYAFQFI------------------------   210
Conservation   ****::***********.:*.***********::*

XCP1          KEDVERVTISGYEDVPENDDESLVKALAHQPVSVAIEASGRDFQFYKGGVFNGKCGTDLD   296
XCP2          KDESETVTINGHQDVPTNDEKSLLKALAHQPLSVAIDASGREFQFYSGGVFDGRCGVDLD   297
NbXCP1        RDESEVVTIDGYHDVPANDEQSLLKALANQPLSVAIEASGRDFQFYSGGVFDGHCGTELD   296
NbXCP2        RDESEVVTIDGYHDVPANDEQSLLKALANQPLSVAIEASGRDFQFYSGGVFDGHCGTELD   296
NbXCP3        -DESEVVTIDGYRDVPANDEQSLLKALANQPVSVAIEASGRDFQFYSGGVFDGHCGSALD   269
Conservation   :: * ***.*:.*** **::**:****:**:****:****:****.****:*:**  **

XCP1          HGVAAVGYGSSKGSDYVIVKNSWGPRWGEKGFIRMKRNTGKPEGLCGINKMASYPTKTK*   355
XCP2          HGVAAVGYGSSKGSDYIIVKNSWGPKWGEKGYIRLKRNTGKPEGLCGINKMASFPTKTK*   356
NbXCP1        HGVAAVGYGTTKGADYIIVKNSWGPKWGEKGFIRMKRNTGKAEGKCGINKMVSFPVKKK*   355
NbXCP2        HGVAAVGYGTTKGADYIIVKNSWGPKWGEKGFIRMKRNTGKAEGKCGINKMVSFPVKKK*   355
NbXCP3        HGVAAVGYGSTKGMDYIIVKNSWGPKWGEKGFIRMKRNTGKPAGICGINKMASFPIKKK*   328
Conservation  *********::** **:********:*****:**:******  * ******.*:* *.**
```

**C**

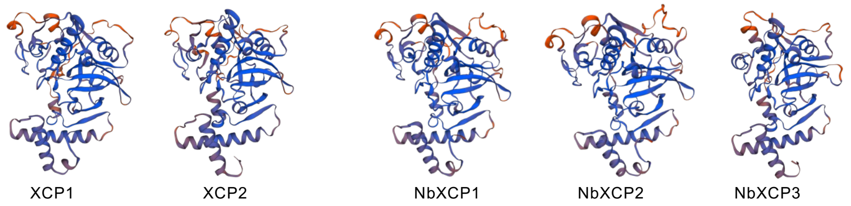


**Supplemental Figure S3** *XCP* homologs in *N. benthamiana*. A, Schematic representation of the NbXCP structures. Sequence information from the *Solanaceae Genomics Network* (https://solgenomics.net/). B, Full-length amino acid sequence alignment for XCPs and NbXCPs (https://www.ebi.ac.uk/Tools/msa/clustalo/). Asterisks represent highly conserved residues. Colons and periods represent low-conserved residues. C, 3D predicted structure for NbXCPs and XCPs (https://swissmodel.expasy.org/).

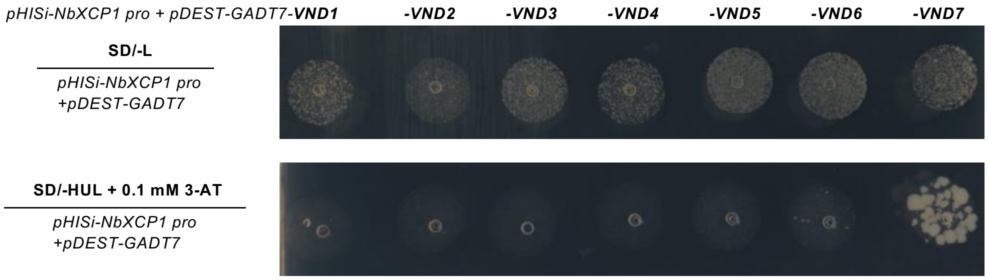


**Supplemental Figure S4** Yeast one-hybrid assay for VNDs binding to *the NbXCP1* promoter. Analysis of yeast transformants harboring *pHISi-NbXCP1 pro* and empty *pDEST-GADT7* vectors or *pHISi-NbXCP1 pro* and *pDEST-GADT7-VNDs* (*VND1–VND7*) vectors on SD (Synthetic Defined) medium. –L: SD medium lacking Leu; –HUL: SD medium lacking His, Ura, and Leu; 3-AT:3-amino-1,2,4-triazole for reducing leaky *HIS* expression.

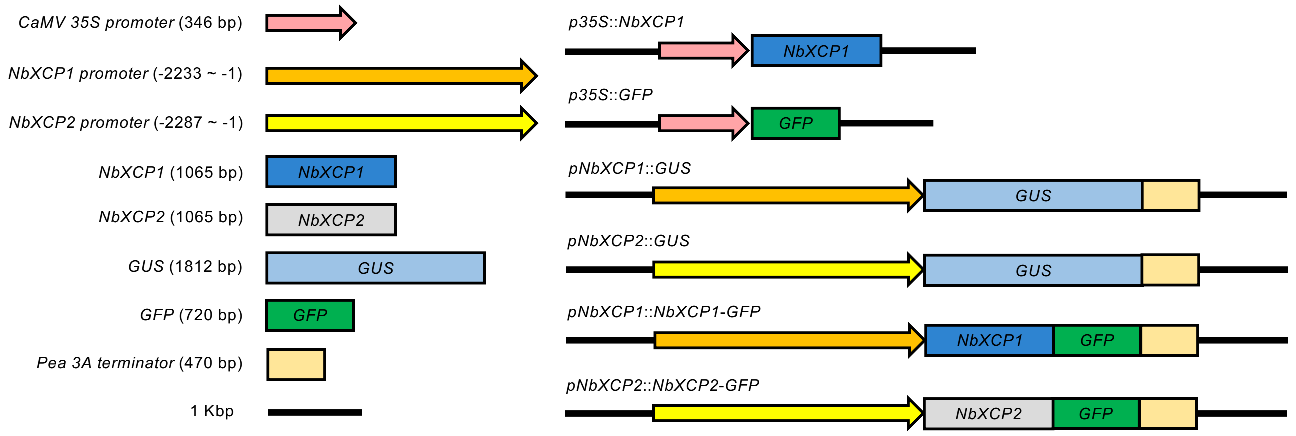


**Supplemental Figure S5** Schematic representation of the GUS and GFP fusion constructs. The upstream sequences (around –1 to –2000 bp) from the first nucleotide of the start codon were selected for the *NbXCP1* and *NbXCP2* putative promoters. The coding sequences of *NbXCP1* and *NbXCP2* were fused to GFP. *pENTR* was an entry vector for recombinational cloning. *pGWB1* was an expression vector for generating transformants.

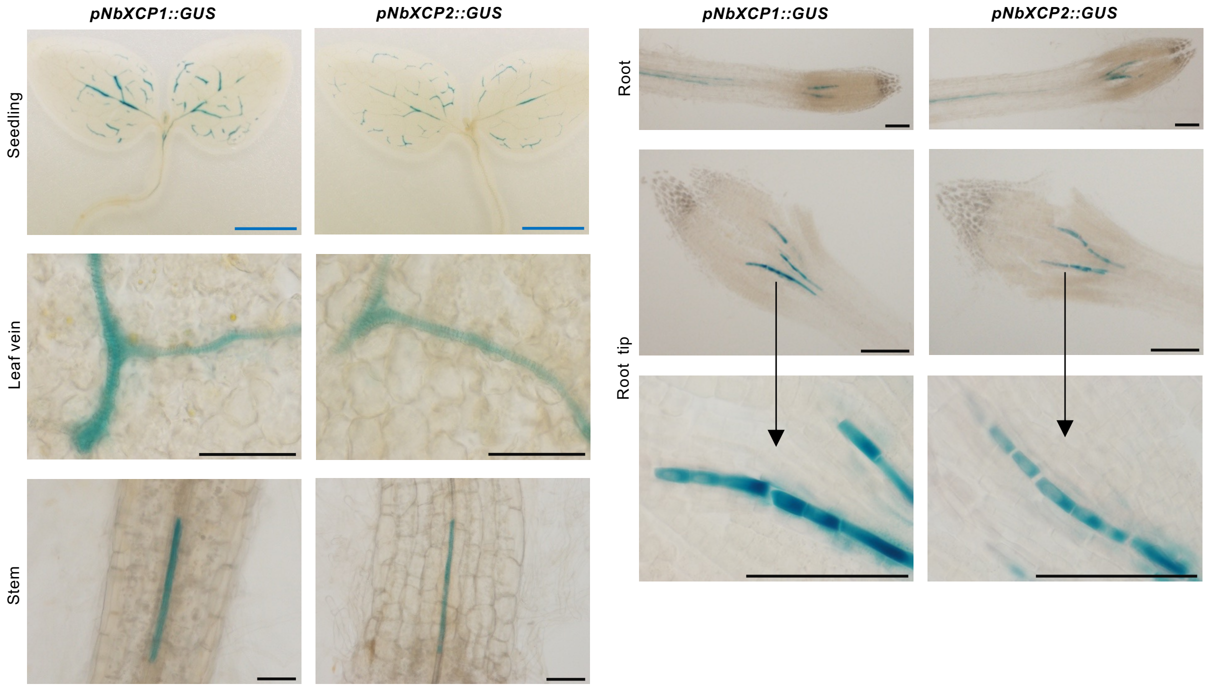


**Supplemental Figure S6** GUS expression analysis of *pNbXCP1::GUS* and *pNbXCP2::GUS* in 7-day-old seedlings. The putative promoters of *NbXCP1* and *NbXCP2* direct GUS expression in the xylem vessels of leaves, stems, and roots. In the root tip, GUS staining was observed in vascular cells without secondary cell walls. Scale bars: blue, 2 mm; black, 100 μm.

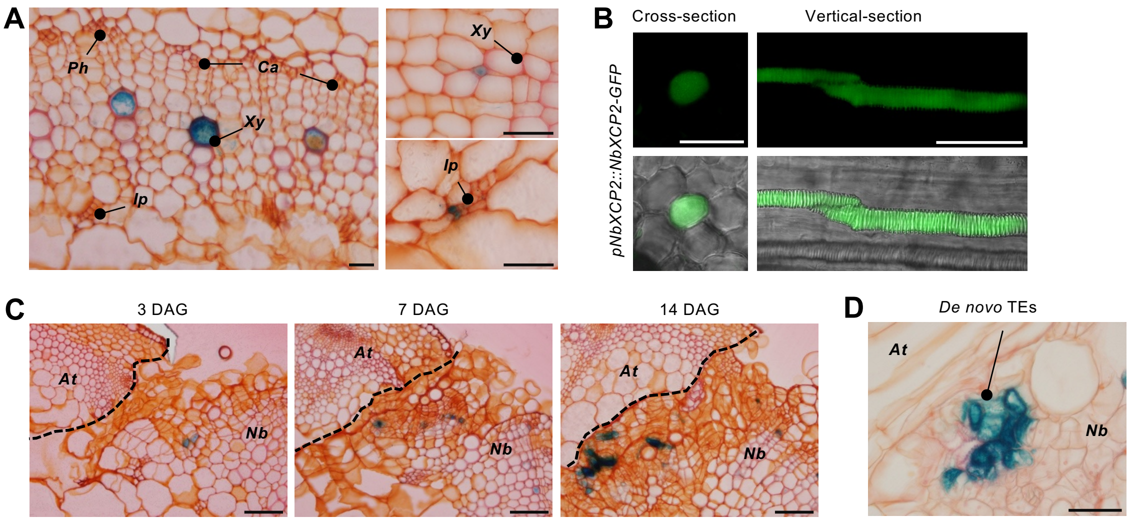


**Supplemental Figure S7** *NbXCP2* expression analysis during graft formation. A, *NbXCP2* promoter directs GUS expression in 4-week-old plant stems. Xylem cells were stained pink with safranin-O. Scale bars, 50 µm. *Ph*: phloem, *Ca*: cambium, *Xy*: xylem, *Ip*: internal phloem. B, GFP expression in xylem vessels. Putative promoters of *NbXCP2* directed NbXCP2-GFP fusion proteins (*pNbXCP2::NbXCP2-GFP*) expression in the cytosol of TEs in 4-week-old plant stems. Scale bars, 100 µm. C, The *NbXCP2* promoter directs GUS expression in the calli of *N. benthamiana* scion at 3, 7, and 14 DAG. The dotted lines indicate the contact surfaces of *N. benthamiana* and Arabidopsis. Scale bars, 100 µm. D, GUS expression in *de novo* TEs. The putative promoters of *NbXCP2* directed GUS expression in *de novo* TEs differentiated in calli at 14 DAG. Scale bars, 50 µm. *At*, Arabidopsis; *Nb*, *N. benthamiana*.

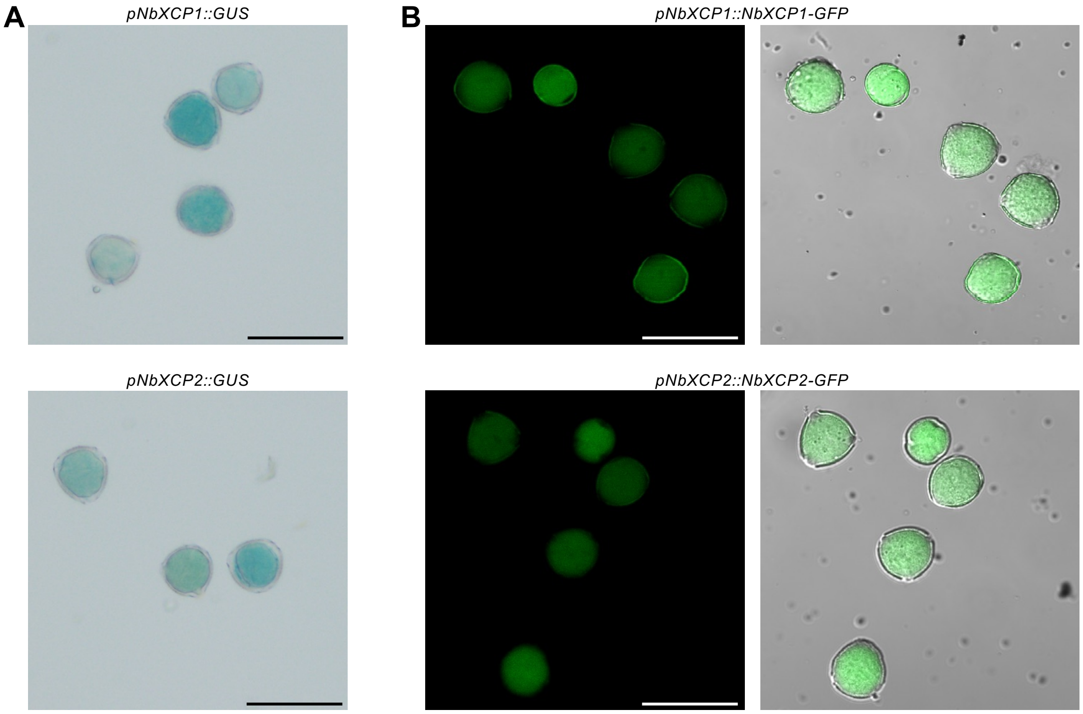


**Supplemental Figure S8** *NbXCP1* and *NbXCP2* expression analysis in the pollen. A, The putative promoters of *NbXCP1* and *NbXCP2* direct GUS (*pNbXCP1::GUS* and *pNbXCP2::GUS*) expression in the pollen. B, The putative promoters of *NbXCP1* and *NbXCP2* directed NbXCP1-GFP and NbXCP2-GFP fusion proteins (*pNbXCP1::NbXCP1-GFP* and *pNbXCP2::NbXCP2-GFP*) expression in the pollen. Scale bars, 50 μm.

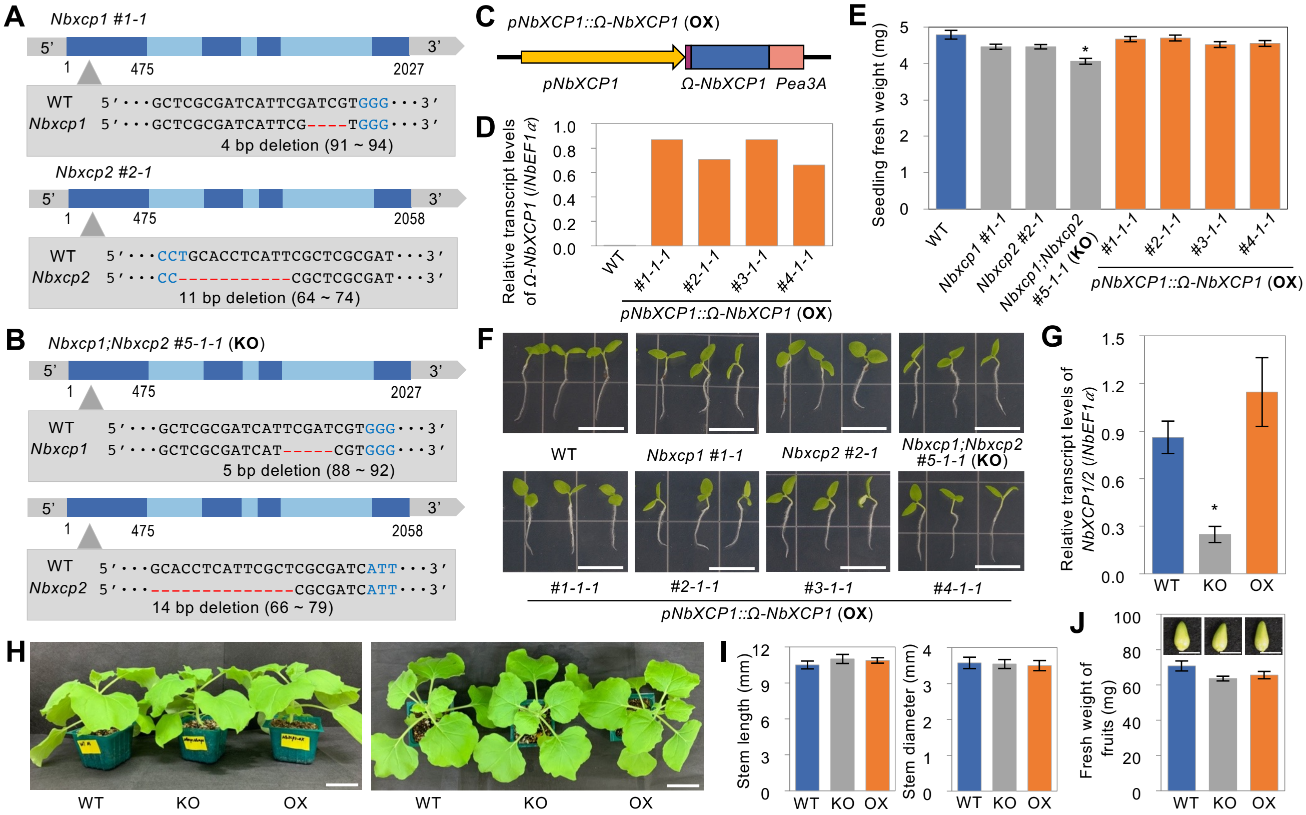


**Supplemental Figure S9** Characterization of *NbXCP* mutants and overexpressors. A, Schematic presentation of CRISPR/Cas9-induced single-knockout mutants (*Nbxcp1* and *Nbxcp2*). B, Schematic presentation of CRISPR/Cas9-induced double-knockout mutant (*Nbxcp1;Nbxcp2*). Dark blue blocks indicate exons, and light blue blocks indicate introns in A and B. C, Structure of the construction for *NbXCP1* translationally enhanced (OX) lines. *pNbXCP1* indicates the 2 kb upstream sequence from the *NbXCP1* start codon. Ω indicates the 5'-leader sequence of tobacco mosaic virus functions as a translational enhancer in plants. *NbXCP1* indicates the CDS sequence of the *NbXCP1* gene. *Pea3A* indicates *Pisum sativum* 3A terminator. D, Transcription of *Ω-NbXCP1* in transgenic plants. qRT-PCR analysis was conducted using 7-day-old seedlings of wild-type (WT) and four OX lines. Fifteen seedlings were pooled and analyzed from each line. E, Fresh weight of 7-day-old wild-type and transgenic seedlings. Error bars indicate mean ± SE (n = 30 for each). Significant differences were determined by multiple comparison analyses according to Tukey's honest significant difference (HSD) test results ($^*P < 0.05$). F, Photographs of wild-type and transgenic *N. benthamiana* lines at 7 days after germination. Scale bars, 1 cm. G, Transcription of endogenous *NbXCP1* and *NbXCP2* in the wild-type and transgenic plants. qRT-PCR analysis was performed using 7-day-old seedlings. Error bars indicate mean ± SE (n = 3). Asterisks represent significant differences determined by Tukey's HSD test ($^*P < 0.05$). H, Representative wild-type and transgenic plants at four weeks after germination. Scale bar, 4 cm. I, Growth of stem before grafting. Stem length and diameter were measured at four weeks after germination. Error bars indicate mean ± SE (n = 32–36 for each). J, Measurement of fruit weight on intact plants at 7-week-old. Scale bars, 5 mm. The fruits were harvested from 6 plants for each transgenic line and wild-type. The measurement with each single fruit. Error bars indicate mean ± SE (n = 15–30 for each). KO indicates homozygous CRISPR/Cas9-induced double-knockout mutant line *Nbxcp1;Nbxcp2 #5-1-1*. OX indicates homozygous *NbXCP1* translationally enhanced line *pNbXCP1::Ω-NbXCP1 #3-1-1* (G–J).

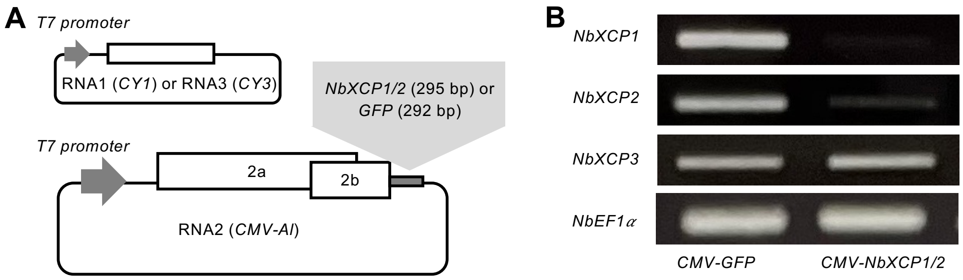


**Supplemental Figure S10** Virus-induced gene silencing (VIGS) for *NbXCP1* and *NbXCP2*. A, Structure of the *cucumber mosaic virus* (*CMV*) vector system. For the VIGS test, a sequence fragment derived from *NbXCP1/2* (the cloned 295 bp sequences are identical between *NbXCP1* and *NbXCP2*) or *GFP* was inserted into the *CMV-Al* vector. The *CY1* and *CY3* vectors generated RNA1 and RNA3 to assist the *CMV*-infection. B, Transcription of *NbXCP* genes in the *CMV*-infected *N. benthamiana* plants at 7 days after infection. RT-PCR analysis was conducted for *NbXCP1–3* using the stem samples. *NbEF1α* was used as a reference. *CMV-GFP* and *CMV-NbXCP1/2* indicate the infected *Nb* plants with *CMV* containing *GFP* and *NbXCP1/2* fragments, respectively.
